# Demonstration of Beat-to-Beat, On-Demand ATP Synthesis in Ventricular Myocytes Reveals Sex-Specific Mitochondrial and Cytosolic Dynamics

**DOI:** 10.1101/2025.07.07.663572

**Authors:** Paula Rhana, Collin Matsumoto, L. Fernando Santana

## Abstract

The energetic demands on ventricular myocytes imposed by the transport of ions and cross-bridge cycling are well known, yet the spatiotemporal dynamics of ATP supply and demand remain poorly understood. Here, using confocal microscopy and genetically encoded fluorescent sensors targeted to mitochondria and cytosol, we visualized beat-to-beat ATP dynamics in ventricular myocytes from male and female mice. These probes showed fluctuations in mitochondrial ATP levels with each contraction, revealing two distinct, spatially localized waveforms—ATP “gain” and ATP “dip”—representing transient increases or decreases in matrix ATP levels, respectively. These waveforms were tightly phase-locked to intracellular Ca^2+^ transients and organized into energetic microdomains. Inhibition of the mitochondrial Ca^2+^ uniporter or the adenine nucleotide translocase attenuated these ATP transients. Although female myocytes exhibited larger mitochondrial ATP transients than their male counterparts, their mitochondrial volume was lower. Female myocytes also exhibited tighter coupling between the sarcoplasmic reticulum and mitochondria and showed a higher density of mitofusin 2 and ATP synthase catalytic α-subunit per unit volume, suggesting more efficient ATP production. Cytosolic ATP transients mirrored mitochondrial waveforms and domain structure in both male and female myocytes. During faster pacing, diastolic cytosolic ATP rose more rapidly in female myocytes, whereas beat-locked ATP transients increased in both sexes but proportionally more in males than in females. These findings demonstrate that ATP is synthesized on a beat-to-beat basis in a modular, microdomain-specific manner. We propose that male myocytes rely on greater mitochondrial mass for energetic scaling, whereas female cells employ architectural precision to optimize ATP delivery.

**Key points summary:**

- It is known that each heartbeat requires precise ATP delivery to fuel ion transport and cross-bridge cycling, but the timing and spatial organization of ATP production in heart cells has been unclear.
- Using advanced imaging and genetically encoded sensors, we visualized beat-to-beat ATP fluctuations in the mitochondria and cytosol of individual male and female mouse ventricular myocytes.
- Mitochondrial ATP levels rose or fell with each beat in spatially confined regions, forming ATP “gain” or “dip” microdomains that were synchronized with Ca^2+^ transients.
- At higher firing rates, beat-locked, diastolic ATP transients rose more quickly in female myocytes, but were larger in male myocytes, highlighting distinct sex-specific strategies for matching energy supply to contractile demand.
- Ventricular myocytes “live paycheck-to-paycheck”, producing just enough ATP on demand to fuel each beat. Male and female myocytes adopt distinct strategies to meet this demand: male myocytes scale output through greater mitochondrial mass, while female myocytes achieve energetic precision via enhanced sarcoplasmic reticulum–mitochondrial coupling.

Abstract figure
Beat-locked mitochondrial ATP transients reveals modular, sex-specific bioenergetic control during excitation-contraction coupling. **(A)** Each action potential activates L-type Ca_V_1.2 channels, producing a Ca^2+^ influx that triggers Ryanodine receptors (RyR2) and elicits SR Ca^2+^ release. **(B)** The cytosolic Ca^2+^ signal is decoded by mitochondria into spatially distinct “high-gain” and “low-gain” regions, shaped by the extent of SR–mitochondrial tethering via mitofusin 2 (Mfn2), yielding heterogeneous mitochondrial activation rather than a uniform, cell-wide metabolic response. **(C)** Mitochondria generate rhythmic, phase-locked ATP transients within discrete microdomains, such that ATP increases and ATP dips can coexist within the same cell. Ca^2+^ entry through the outer membrane (via VDAC) and into the matrix (via MCU) stimulates oxidative phosphorylation, increasing ATP production; ATP is exported by ANT to create local cytosolic “supply bursts” aligned with beat-to-beat demand (“paycheck-to-paycheck” energetics). Female myocytes show a higher prevalence of tightly coupled, high-gain ATP-producing microdomains, whereas male myocytes display a shifted balance toward lower-gain regions, consistent with sex-dependent SR–mitochondrial coupling and ATP microdomain patterning.

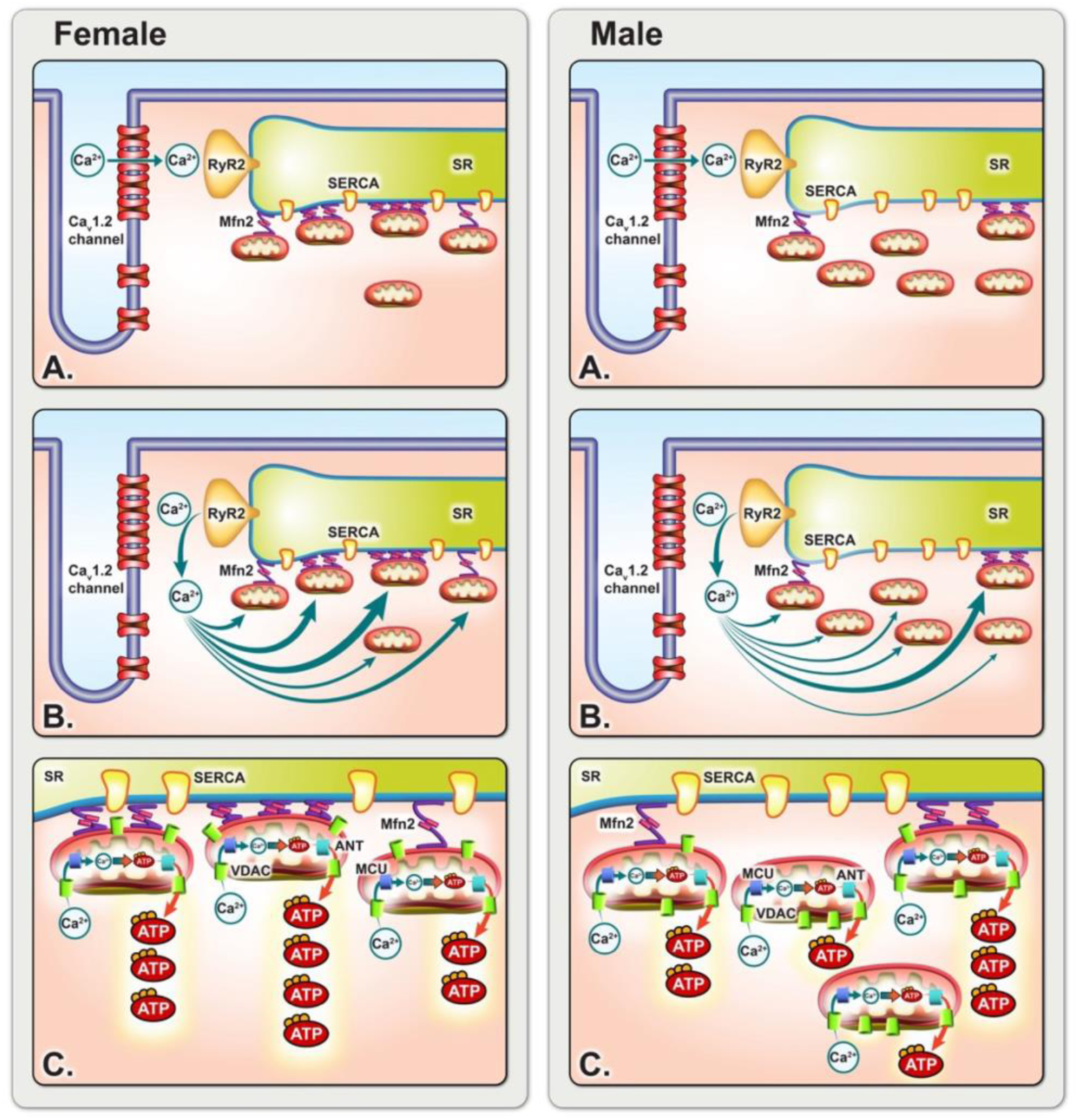

## Introduction

The function of the heart is to pump blood through systemic and pulmonary circulations, delivering oxygen and nutrients to tissues and removing metabolic waste. To accomplish this, the heart contracts rhythmically—about 100,000 times per day in humans and 1 million times in mice—through a sequence of electrical and mechanical events known as the cardiac cycle. Each cycle begins with the generation of an action potential by pacemaker cells in the sinoatrial node and culminates in ventricular contraction and relaxation—tightly regulated processes that ensure adequate cardiac output under varying physiological demands.

The conversion of electrical excitation into mechanical force is mediated by a process termed excitation–contraction (EC) coupling (Eisner *et al*., 2017). In ventricular myocytes, membrane depolarization triggers the opening of clusters of L-type Ca^2+^ channels (Ca_V_1.2) in transverse tubules (Dixon *et al*., 2012; Dixon *et al*., 2015; Westhoff *et al*., 2024; Spooner *et al*., 2025), allowing a small influx of Ca^2+^ into the cytosol (Santana *et al*., 1996; Wang *et al*., 2001). This local Ca^2+^ signal activates ryanodine receptors in the adjacent junctional sarcoplasmic reticulum (SR) through a Ca^2+^-induced Ca^2+^-release mechanism (Fabiato, 1983). The resulting “Ca^2+^ sparks” raise intracellular cytosolic Ca^2+^ concentration ([Ca^2+^]_i_) and initiate actomyosin cross-bridge cycling and myocyte contraction (Cheng *et al*., 1993; Cannell *et al*., 1995; López-López *et al*., 1995). This is followed by relaxation as Ca^2+^ is re-sequestered into the SR by the sarcoplasmic/endoplasmic reticulum Ca^2+^-ATPase (SERCA) pump and extruded from the cell by the Na^+^/Ca^2+^ exchanger, allowing the cycle to repeat (Balke *et al*., 1994; Li *et al*., 1998).

ATP fuels SERCA-driven Ca^2+^ reuptake, ion transport (via pumps and exchangers), and actomyosin cross-bridge cycling, with ∼95% of this ATP supplied by mitochondrial oxidative metabolism and the remaining ∼5% by glycolysis (Wisneski *et al*., 1990; Saddik & Lopaschuk, 1991; Zhou & Tian, 2018; Lopaschuk *et al*., 2021). Classic work has established coupling between Ca^2+^ handling and ATP production, showing that Ca^2+^ entering the mitochondrial matrix via the mitochondrial Ca^2+^ uniporter (MCU) activates three Ca^2+^-sensitive dehydrogenases—pyruvate, isocitrate, and 2-oxoglutarate dehydrogenase; this boosts NADH production and drives proton flux through the ATP5A-containing F_1_F_o_-ATP synthase (Denton *et al*., 1980; Denton & McCormack, 1990; McCormack *et al*., 1990; Kirichok *et al*., 2004; Garbincius & Elrod, 2022). In this process, cytosolic ADP and P_i_ must access the matrix and newly synthesized ATP must be exported, a transfer mediated by voltage-dependent anion channels (VDAC) in the outer mitochondrial membrane and the adenine nucleotide translocase (ANT) in the inner membrane. Accordingly, mitochondrial efficiency is shaped not only by enzymatic capacity but also by membrane transport, organelle architecture, and proximity to SR Ca^2+^-release sites. The tethering protein mitofusin-2 (Mfn2), located on the outer mitochondrial membrane, mediates physical and functional coupling between mitochondria and the junctional SR, facilitating rapid Ca^2+^ transfer and enhancing oxidative phosphorylation (Chen *et al*., 2012; Dorn *et al*., 2015). Mitochondrial ATP production is supplemented by glycolysis and is further supported by creatine and adenylate kinase reactions, which draw on phosphocreatine and ADP pools to regenerate ATP. These pathways act in concert to maintain ATP supply across varying workloads. Despite the substantial energetic cost of each contraction cycle, prevailing models have assumed that intracellular ATP remains effectively constant due to the high throughput of mitochondrial ATP production, the rapid kinetics of phosphotransfer reactions, and the supplementary contribution of glycolysis (Chen *et al*., 1998; Murphy & Steenbergen, 2008).

High-resolution imaging of intracellular ATP concentration ([ATP]_i_) in beating ventricular myocytes has now upended this view. In Rhana *et al*. (2024), we showed that cytosolic ATP fluctuates with each ventricular beat and that these oscillations disappear when oxidative phosphorylation is inhibited, revealing the limited buffering capacity of phosphotransfer and the dominant role of mitochondria in sustaining EC coupling. These findings raise two questions that are addressed in the present study: (i) How tightly are SR Ca^2+^ release and mitochondrial ATP synthesis temporally linked during each beat? And (ii), given emerging evidence for sex-specific differences in mitochondrial content and function (Cao et al., 2022), how do the kinetics and amplitudes of cytosolic and mitochondrial ATP transients differ between male and female cardiomyocytes?

In this study, we address these questions by directly monitoring cytosolic and mitochondrial ATP dynamics in beating male and female ventricular myocytes using newly developed expressible indicators, revealing a previously unrecognized, modular and beat-synchronized mode of ATP production. By comparing mitochondrial and cytosolic ATP fluctuations within the same cell type under physiological pacing conditions, we test the hypothesis that mitochondrial ATP, like its cytosolic counterpart, varies beat to beat in response to EC coupling. Building on evidence that mitochondrial volume, gene expression, and SR–mitochondrial coupling differ between male and female hearts (Cao *et al*., 2022; Clements *et al*., 2023), we further examine how stimulation frequency and biological sex shape these dynamics.

Our results reveal that cytosolic and mitochondrial ATP oscillate in synchrony with the cardiac cycle and show that these oscillations are modulated by stimulation frequency and differ according to sex. Female myocytes exhibit a faster rise in diastolic cytosolic ATP during increased pacing that coincides with stronger SR–mitochondrial coupling and higher expression of Mfn2 and ATP5A. In contrast, mitochondrial volume is greater in male myocytes, and the amplitude of beat-to-beat ATP transients in these cells is selectively increased under load. Together, these findings support a model in which mitochondria participate dynamically in the beat-to-beat regulation of cardiac energetics and suggest sex-specific strategies—mass-based scaling versus architectural precision—for matching energy supply to contractile demand.

## Methods and Materials

### Ethical approval

This study adhered to the guidelines outlined in the National Institutes of Health (NIH) Guide for the Care and Use of Laboratory Animals. The Institutional Animal Use and Care Committee (IACUC) at the University of California Davis approved the animal use and protocols described (IACUC #24065), which align with ARRIVE guidelines.

### Animal source and housing

Adult male and female C57BL/6J mice (8–12 weeks old) were obtained from The Jackson Laboratory (USA). Animals were housed in standard cages on a 50:50 light/dark cycle with unrestricted access to food and water *ad libitum*.

### Infection of mice with adeno-associated viruses (AAVs)

Adeno-associated virus serotype 9 (AAV9) particles carrying either the cytosolic iATP reporter iATPSnFR^1.0^ (cyto-iATP) (Lobas *et al*., 2019) or a mitochondria-targeted iATP reporter iATPSnFR^2.0^ (mito-iATP) (Marvin *et al*., 2024) were prepared at 4 × 10^12^ viral genome copies per milliliter (vg/mL). Mice were induced with 5% isoflurane in oxygen (0.8–1.2 L/min) and maintained at 2% via a nose cone. Depth of anesthesia was confirmed before the procedure and monitored throughout the procedure by the absence of the pedal withdrawal response to a firm toe pinch. For retro-orbital intravenous delivery, 100 µL of viral suspension was administered using a sterile insulin-type micro syringe and one drop of ophthalmic lubricant was applied after the injection. Post-procedure checks were done after 1 h and at least once daily for 24–48 h.

### Isolation of mouse ventricular myocytes

Mice were euthanized 1–2 weeks post AAV infection and cardiomyocyte isolation was performed as previously described (Shioya, 2007; Rossow *et al*., 2009). Briefly, mice were heparinized (1000 UI/Kg, i.p.) and a lethal dose of pentobarbital (100 mg/kg, i.p.) was administered. Hearts were rapidly excised and plunged into cold cell isolation buffer (130 mM NaCl, 5 mM KCl, 0.5 mM MgCl_2_, 0.33 mM NaH_2_PO_4_, 25 mM HEPES, 22 mM glucose, and 3mM C_3_H_3_NaO_3_, pH 7.4 adjusted using NaOH) supplemented with 150 µM EGTA. After aortic cannulation, an enzyme-containing cell isolation buffer (0.04 mg/mL protease type 2 collagenase, supplemented with 50 µM CaCl_2_) at 37 _o_C was retrogradely perfused for 6–9 min until flow increased. Ventricles were removed and minced for further digestion in fresh enzyme solution. The suspension was filtered and Ca^2+^ restored stepwise. Isolated ventricular myocytes were maintained at room temperature (∼24°C) in Tyrode’s solution (140 mM NaCl, 5.4 mM KCl, 1.8 mM CaCl_2_, 1 mM MgCl_2_, 5 mM HEPES, and 5.5 mM glucose, pH 7.4 adjusted using NaOH) and used within 5 hours of isolation. To assess the role of other mitochondrial substrates in ATP dynamics, we perfused myocytes with a “physiological mix” of Tyrode’s containing 0.1 mM C_3_H_3_NaO_3_, 1 mM C_3_H_5_NaO_3_, 2 mM L-Carnitine, 0.3 mM Palmitate (BSA complex solution).

### Field stimulation

Action potentials were evoked by field stimulation using two platinum wires separated by 0.5 cm, positioned at the bottom of the perfusion chamber. Square, 4-ms voltage pulses were generated using a Grass stimulator (AstroMed Inc., USA). Pulse amplitudes were 10–40 V and delivered at frequencies ranging from 1-2 Hz.

### Confocal imaging of action potential-evoked [Ca^2+^]_i_, [ATP]_i_, and [ATP]_mito_ signals

Cytosolic Ca^2+^ was imaged by loading ventricular cardiomyocytes expressing the cyto-iATP or mito-iATP sensor with 10 µM Rhod-3-AM (Thermo Fisher, USA) following manufacturer’s guidelines. After loading with Rhod-3, a drop of the cell suspension was transferred to a temperature-controlled (37°C) perfusion chamber on a microscope stage and allowed to settle onto the coverslip for 5 minutes. Cells were then perfused with Tyrode’s solution and field stimulated. Diastolic and systolic [Ca^2+^]_i_ and [ATP]_i_ or [ATP]_mito_ signals were simultaneously recorded using an inverted Olympus FV3000 confocal microscope (Olympus, Japan) operating in 2D or line-scan mode. Fluorescent indicators were excited with solid-state lasers emitting at 488 nm (cyto-iATP and mito-iATP) or 561 nm (Rhod-3) through a 60× oil-immersion lens (PlanApo) with a numerical aperture (NA) of 1.40. During analysis, background was subtracted from all confocal images, and fluorescence signals were expressed as F/F₀, where F is the fluorescence intensity at a given time point, and F_0_ is the mean baseline fluorescence. Cyto-iATP fluorescence values were converted to concentration units using the pseudo-ratiometric method (Cheng *et al*., 1993). A dissociation constant (K_d_) of 1460 µM and a resting level of [ATP]_i_ of 457 µM was used in these calculations (Rhana *et al*., 2024).

### Ratio-metric mito-iATP/HaloTag recordings

Isolated myocytes co-expressing the mito-iATP sensor and HaloTag were incubated for 30 min at 37 °C with the HaloTag ligand JFX554 (Promega, USA; #HT1030) at a final concentration of 0.1 µM. After rapid solution exchange, cells were transferred to pre-warmed (37 °C) Tyrode’s solution and placed in a perfusion chamber on the microscope stage. Myocytes were allowed to settle onto the coverslip for 5 min before imaging. Images were acquired as described above (*Confocal imaging* section) using solid-state laser excitation at 488 nm (mito-iATP) and 561 nm (HaloTag).

### Super-resolution radial fluctuation (SRRF) imaging

Freshly isolated ventricular myocytes expressing cyto-iATP or mito-iATP sensors were incubated for 30 minutes at 37°C with 250 nM MitoTracker Deep Red (Molecular Probes, USA). Two-dimensional SRRF images were acquired using Fusion software on an Andor Dragonfly 200 spinning-disk confocal platform (Andor Technologies, UK) coupled to an inverted Leica DMi* microscope fitted with a 60× oil-immersion objective (NA = 1.40) and Andor iXon EMCCD camera. Colocalization analyses for mito-iATP versus MitoTracker and cyto-iATP versus MitoTracker were performed using IMARIS software (colocalization tool).

### VDAC1–SERCA2 colocalization analysis

Immunofluorescence labeling was performed on freshly dissociated ventricular myocytes (wild-type, non-AAV infected adult C57BL/6J mice). Myocytes were left to adhere for 1 hour at room temperature prior to fixation, in coverslips coated with Poly-L-lysine and Laminin. Myocytes were fixed with 4% formaldehyde diluted in phosphate-buffered saline (PBS) (Fisher Scientific, USA) for 15 min at room temperature, washed, and incubated with 50 mM glycine for 10 min to reduce aldehydes. Cells were then incubated in blocking buffer made of 3% w/v bovine serum albumin and 0.25% Triton X-100 in PBS, followed by incubation with mouse anti-SERCA2 (#MA3-919, Invitrogen, USA; 1:100) and mouse anti-VDAC1 (#ab14734, Abcam, USA; 1:100) diluted in blocking buffer overnight at 4 °C. Myocytes were washed, incubated at room temperature for one hour with Alexa Fluor 488-conjugated goat anti-mouse IgG2b (#A21141, Invitrogen, USA; 1:1000) and Alexa Fluor 647-conjugated goat anti-mouse IgG2a (#A21241, Invitrogen, USA; 1:1000) diluted in blocking buffer followed by washes in PBS. All washes were performed with PBS three times for 5 minutes. Coverslips were mounted onto microscope slides in Vectashield mounting medium (Vector Labs, USA) and sealed with clear nail polish. Images were collected on a Dragonfly 200 spinning disk confocal (Andor, UK), coupled to a DMi* Leica microscope (Leica, Germany) equipped with a 60x oil immersion objective (NA = 1.40) and acquired using an Andor iXon EMCCD camera. Two dimensional wild-field images were collected via Fusion software. Using ImageJ, images were background-subtracted, and each channel was subject to thresholding to generate binary masks and image math was performed to visualize and quantify the SERCA2-VDAC1 colocalization.

### Mitochondrial volume analysis

Ventricular myocytes from male and female mice were incubated with 250 nM MitoTracker Deep Red (Thermo Fisher Scientific, USA) for 30 minutes at 37°C. Cells were allowed to adhere to poly-L-lysine/laminin-coated coverslips for 1 hour at room temperature, then fixed with 4% formaldehyde diluted in PBS for 15 minutes. Coverslips were mounted using VectaShield mounting medium and sealed with clear nail polish. Confocal z-stacks were acquired on an Olympus FV3000 confocal microscope using a 60× oil-immersion objective (NA = 1.40), with a z-step of 0.5 µm. After importing three-dimensional (3D) image stacks into IMARIS 10 (Andor, UK), mitochondria were segmented based on MitoTracker fluorescence intensity using the Surfaces tool and a consistent thresholding approach. Total mitochondrial volume was quantified for each cell and normalized to cytosolic volume to obtain the mitochondrial volume fraction.

### Western blots

Female and male ventricles were collected and homogenized in ice-cold RIPA buffer supplemented with a cocktail of protease inhibitors (Pierce Protease Inhibitor Mini Tablets; Thermo Scientific, USA). Proteins (60 μg) were separated by sodium dodecyl sulfate-polyacrylamide gel electrophoresis (SDS-PAGE) followed by semi-dry transfer onto a PVDF membrane (Merck, USA). Membranes were washed in Tris-buffered saline containing 0.1% Tween-20 (TBS-T), blocked in 5% bovine serum albumin (BSA) TBS-T for 1 hour at room temperature, and incubated with rabbit anti-Mfn2 (#9482S, Cell Signaling, USA; 1:500), mouse anti-ATP5A1 (#459240, Invitrogen, USA; 1:1000) and mouse anti-VDAC1 (clone N152B/23, RRID 2877354; UC Davis/NIH Neuromab Facility, USA; 1:200) diluted in blocking buffer. Membranes were then rinsed with TBS-T and incubated with mouse IgG 800CW or rabbit IgG 680CW secondary antibodies (1:15,000; Li-Cor, USA) for 1 hour at room temperature. Fluorescence signals were detected using an Odyssey Infrared Imager (LiCor, USA). Blots were digitized, and bands quantified using ImageJ. Protein levels were expressed as the ratio of optical densities of specific bands to VDAC1.

### Statistics

Data are presented as mean ± SD, from *n* cells, sites or transients, and *N* animals. Normality was assessed for all data sets (Shapiro–Wilk test) and passed it (P > 0.05). Accordingly, only parametric statistics were implemented. Hierarchical statistics (nested *t*-tests) and paired Student’s *t*-tests were used throughout the paper, as indicated in the figure legend. Exact P values are provided in the text and/or figures, with P < 0.05 considered statistically significant. All data points are shown in the figures, with light symbols representing individual cells and dark symbols individual animals.

## Results

### Mitochondrial ATP transients exhibit two beat-locked waveforms with a sex-dependent spatial distribution

Mitochondrial ATP was imaged using a mitochondrial matrix-targeted intracellular ATP (iATP) sensor termed mito-iATP, a version of the low-affinity A95A/A119L variant (apparent K_d_ ≈ 0.5 mM) of the high-dynamic-range reporter, iATPSnFR2, created by fusing four tandem COX8 leader sequences to the N-terminus of iATPSnFR2 and cloning it into an AAV9 cassette, as described by Marvin *et al*. (2024). Super-resolution radial fluctuation (SRRF) imaging performed on adult mouse ventricular myocytes expressing mito-iATP (**Figure 1A**) showed that mito-iATP fluorescence formed a finely striated reticulum that colocalized with MitoTracker Deep Red, with no detectable cytosolic signal in either male or female myocytes. A magnified 10 x 10 µm view (**Figure 1A insets)** illustrates the pixel-level concordance between the two channels. A quantitative colocalization analysis yielded Manders’ coefficients of 0.94 ± 0.02 for female and 0.98 ± 0.02 for male myocytes, indicating precise targeting throughout the mitochondrial lattice. As shown in **Figure 1B**, brief bath application of 1 µM FCCP, an uncoupler of oxidative phosphorylation, decreased mito-iATP fluorescence from a normalized F/F_0_ value of 1.00 to 0.59 ± 0.13 in female (*P* < 0.0001) and 0.55 ± 0.12 in male (*P* = 0.0002), demonstrating that mito-iATP reliably reports dynamic changes in matrix ATP concentration. No difference was observed in the FCCP-induced minimum fluorescence between female and male myocytes (P = 0.7748).

**Figure 1.**
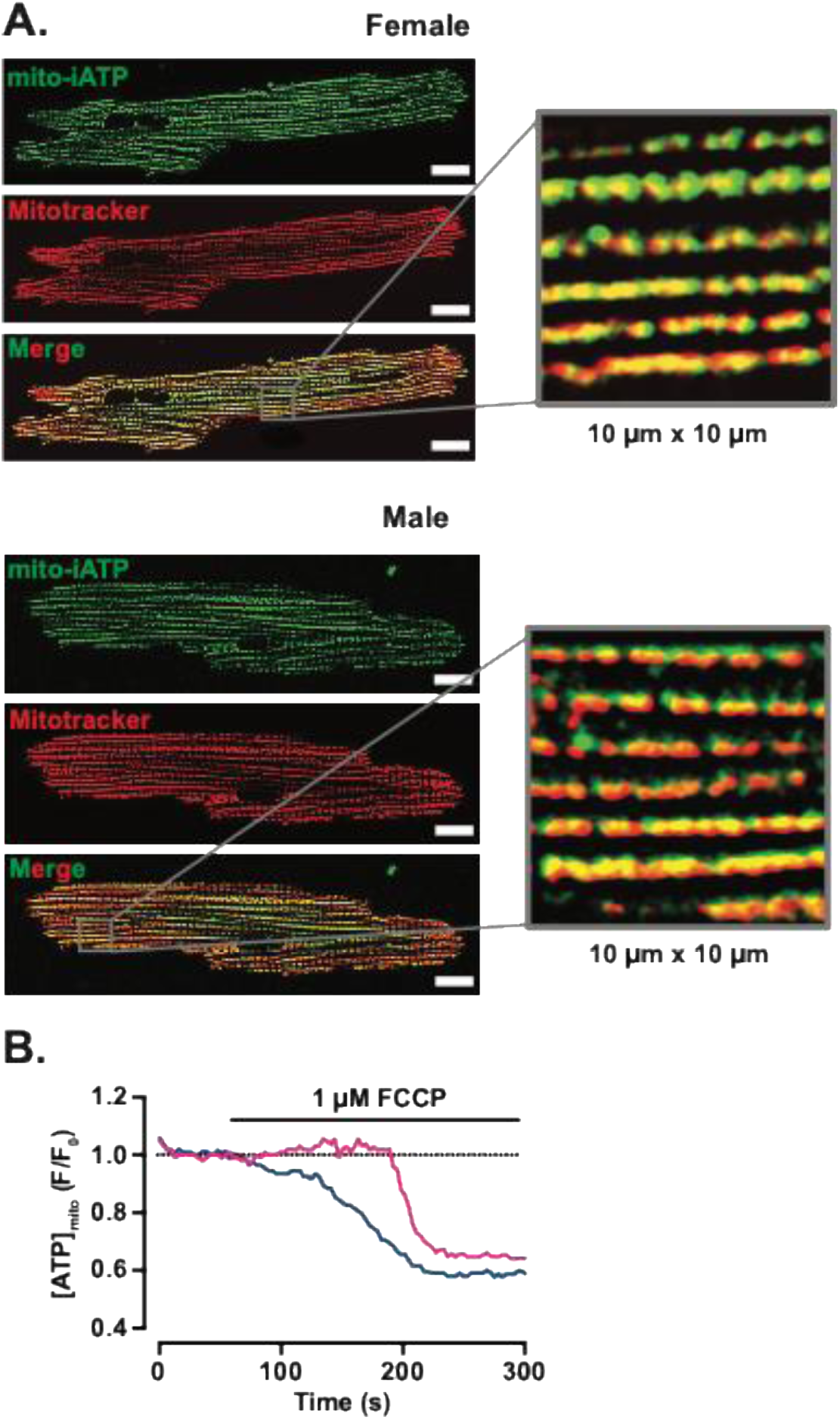
Targeting and validation of the mitochondrial ATP sensor, mito-iATP, in adult mouse ventricular myocytes. **(A)** Live-cell, wide-field confocal images of myocytes expressing mito-iATP (green) and Mitotracker Deep Red (red). The merged panel and 10 x 10 µm inset demonstrate pixel-level concordance between channels. Manders colocalization coefficient: female, 0.94 ± 0.02 (N = 3 mice, *n* = 26 cells); male, 0.98 ± 0.02 (N = 3 mice, *n* = 33 cells). Data are presented as mean ± SD. Scale bars: 10 µm. **(B)** Time course of normalized mito-iATP fluorescence (F/F_0_) after bath application of 1 µM FCCP (female: N = 3 mice, *n* = 10 cells; male: N = 2 mice, *n* = 11 cells).

To test whether [ATP]_mito_ fluctuates on a beat-to-beat basis, we loaded myocytes expressing mito-iATP with the red-shifted Ca^2+^ indicator Rhod-3 and applied field stimulation at 1 Hz. We chose 1 Hz as a standard baseline pacing rate for isolated adult ventricular myocytes because it provides stable Ca^2+^ cycling and preserves cell health while allowing accurate temporal alignment of Ca^2+^ and ATP signals with minimal rundown. Although mouse hearts beat faster in vivo, the core Ca^2+^-dependent control of mitochondrial metabolism is conserved; thus, 1 Hz provides a well-controlled reference condition for mechanistic experiments, with frequency-dependent behavior examined separately at higher pacing rates.

Representative line-scan images (**Figure 2A, B**) captured simultaneous [Ca^2+^]_i_ (upper trace, red) and matrix ATP (lower trace, green) responses during individual action potentials. Each stimulus elicited a phasic [ATP]_mito_ fluctuation that was tightly phase-locked to the [Ca^2+^]_i_ transient. Two reproducible waveform types were identified. In the first (Mode 1), [ATP]_mito_ rose rapidly and then returned to baseline (**Figure 2A**). In the second (Mode 2), [ATP]_mito_ dipped transiently and recovered with similar kinetics (**Figure 2B**). These opposite-polarity responses coexisted within single myocytes, but were spatially segregated, revealing a patchwork of energetic microdomains. Importantly, the amplitude of whole-cell [Ca^2+^]_i_ transients was similar in cells displaying either Mode 1 or Mode 2 fluctuations across both sexes (**Figure 2C**) (2.40 ± 0.53 and 2.16 ± 0.5 F/F_0_ for Mode 1 sites in female and male, respectively - P = 0.2289; and 2.59 ± 0.56 and 2.58 ± 1.07 F/F_0_ for Mode 2 sites in female and male, respectively – P = 0.8580). The average [ATP]_mito_ amplitudes of Mode 1 sites (1.53 ± 0.27 F/F_0_ in females and 1.46 ± 0.20 F/F_0_ in males, P = 0.4124) and Mode 2 sites (0.71 ± 0.08 F/F_0_ in females and 0.70 ± 0.09 F/F_0_ in males, P = 0.5883) were similar across sexes (**Figure 2D**).

**Figure 2.**
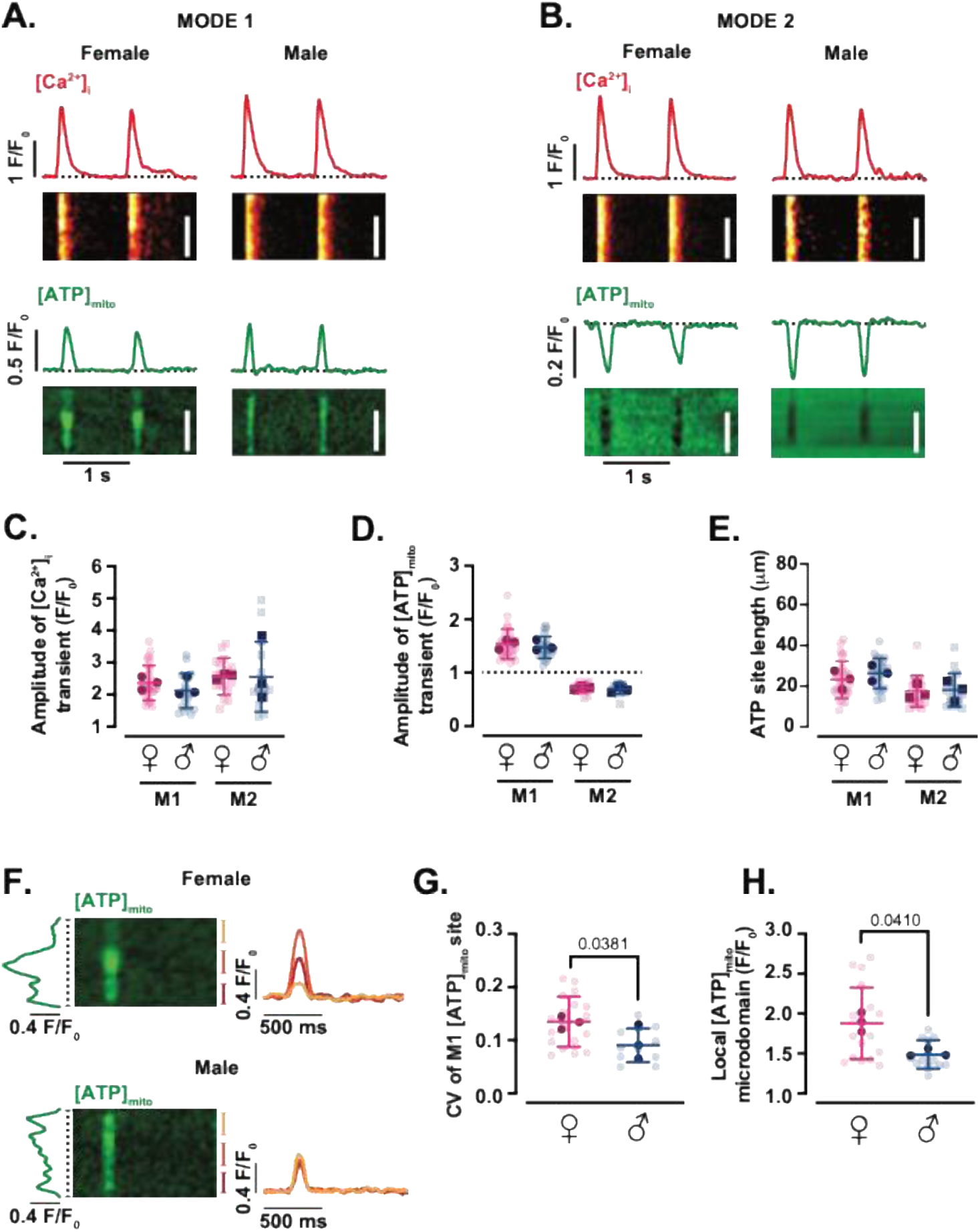
Beat-locked mitochondrial ATP transients adopt two discrete waveforms. (Modes 1 and 2). Representative line-scan images of [Ca^2+^]_i_ (top) and [ATP]_mito_ (bottom) from female and male myocytes with Mode 1 **(A)** or Mode 2 **(B)** ATP dynamics. Red and green traces above each line-scan show [Ca^2+^]_i_ and [ATP]_mito_ time courses, respectively. Scale bars: 20 µm. Scatter plots of the amplitudes of [Ca^2+^]_i_ **(C)** and [ATP]_mito_ **(D)** transients, and spatial spread of [ATP]_mito_ **(E)** in female (magenta) and male (blue) myocytes (female: N = 3 mice, *n* = 27 and 17 cells of Mode 1 and 2 sites, respectively; male: N = 3 mice, *n* = 21 and 15 cells of Mode 1 and 2 sites, respectively). **(F)** Representative regions of interest in Mode 1 [ATP]_mito_ microdomains in female (top) and male (bottom) myocytes. Green trace (left) is the spatial profile of the [ATP]_mito_ signal oriented along the dotted line. Red, yellow, and orange traces are normalized fluorescence intensity of the local [ATP]_mito_ microdomain. Scatter plots of the coefficient of variation (CV) of Mode 1 [ATP]_mito_ sites **(G)** and peak amplitude for the [ATP]_mito_ microdomain **(H)** (female: N = 3 mice, *n* = 18 cells; male: N = 3 mice, *n* = 12 cells). Data are presented as mean ± SD. All significant values are provided from a nested *t* test. Light symbols: individual cells; dark symbols: individual animals.

Quantification of over 30 individual mito-iATP sites revealed a modest, but consistent, sex difference in waveform prevalence. In females, 43% of cells exhibited Mode 1 exclusively, 7% exhibited Mode 2 exclusively, and 50% showed both modes; in males, the corresponding proportions were 38%, 8%, and 54%. Despite this difference in spatial distribution, the average length of local transients was indistinguishable between sexes: Mode 1 sites averaged around 25 µm (23 ± 9 µm in females and 26 ± 7 µm in males, P = 0.4279), whereas Mode 2 sites averaged 18 ± 8 µm in both sexes (P = 0.8578) (**Figure 2E**). Given that a single mitochondrion spans ∼1.5–2 µm, these measurements suggest that each narrow line-scan region captures synchronous ATP fluctuations across strings of adjacent ∼10–18 interfibrillar mitochondria (Moreland, 1962; Hom & Sheu, 2009), confirming the microdomain nature of these events.

Despite comparable amplitudes, the spatial patterning of [ATP]_mito_ elevations along interfibrillar mitochondrial strings was more heterogeneous in females (**Figure 2F**). To quantify this, we calculated the coefficient of variation (CV = standard deviation/mean) of mito-iATP signals along the transient peak. The CV was significantly greater in female (0.14 ± 0.05) than male (0.09 ± 0.03) myocytes (*P* = 0.0381) (**Figure 2G**), suggesting greater spatial variability in mitochondrial ATP synthesis. Consistent with this, ATP “hotspots” in cardiomyocytes reached higher peak values in females (1.91 ± 0.46 F/F_0_) than males (1.51 ± 0.18 F/F_0_; *P* = 0.0410), even though the average amplitude per site remained equivalent (**Figure 2H**).

Because standard Tyrode’s solution contains glucose as the only exogenous carbon source, we asked whether supplementing the bath with a more physiological mixture of metabolic substrates would alter beat-locked ATP dynamics (**Figure 3**). To approximate the metabolic milieu of arterial blood, we perfused myocytes with a “physiological mix” containing glucose together with a cocktail of long-chain fatty acids (palmitate) and additional oxidizable substrates (lactate, pyruvate and L-carnitine), with BSA serving as a carrier for lipid species. Under these conditions, the amplitude of mito-iATP transients in both Mode 1 (ATP-gain) and Mode 2 (ATP-dip) microdomains were indistinguishable from those recorded in Tyrode’s solution (**Figure 3B**) (Mode 1: 1.62 ± 0.23 F/F_0_ in female, P = 0.2625 vs control; and 1.62 ± 0.18 F/F_0_ in male, P = 0.1003 vs control; Mode 2: 0.66 ± 0.09 F/F_0_ in female, P = 0.1597 vs control; and 0.64 ± 0.09 F/F_0_ in male, P = 0.2510 vs control). Likewise, [Ca^2+^]_i_ transient amplitude were similar in Tyrode and physiological mix (**Figure 3C**) (Mode 1: 2.41 ± 0.31 F/F_0_ in female, P = 0.9548 vs control; and 2.36 ± 0.45 F/F_0_ in male, P = 0.1255 vs control; Mode 2: 2.41 ± 0.35 F/F_0_ in female, P = 0.3068 vs control; and 2.33 ± 0.56 F/F_0_ in male, P = 0.7855 vs control). Thus, ATP transients are preserved when myocytes are supplied with a physiological substrate mix, indicating that these signals are not an artifact of perfusion with a simplified glucose-only Tyrode solution.

**Figure 3.**
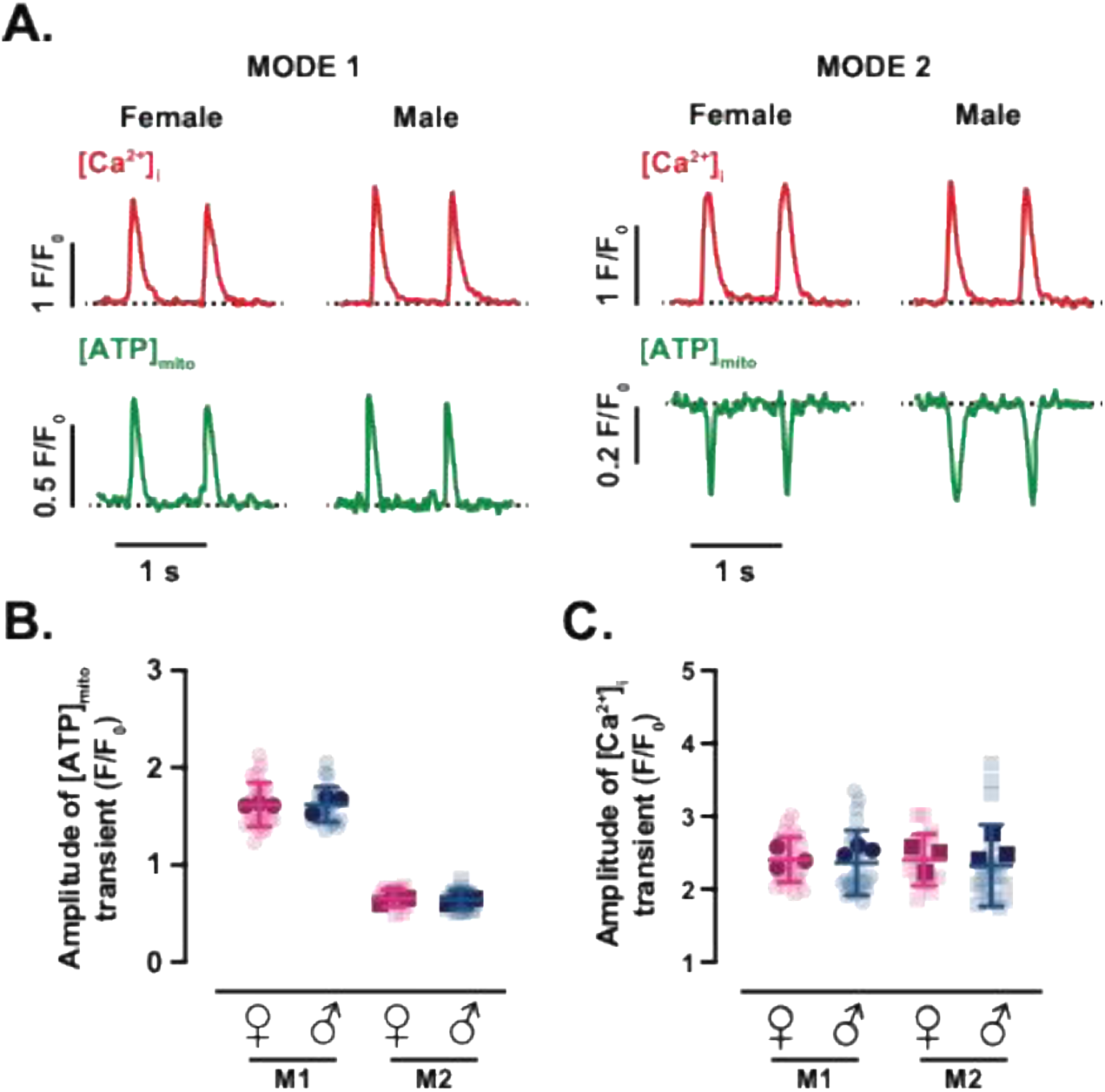
Effect of mixed substrates on mitochondrial ATP and [Ca²⁺]_i_ transient measurements. **(A)** Representative time course from line scans of [Ca^2+^]_i_ (top) and [ATP]_mito_ (bottom) in female and male myocytes with Mode 1 (left) or Mode 2 (right) ATP dynamics using a new physiological substrate mix containing pyruvate, lactate, carnitine, and palmitate. Scatter plots of the amplitudes of [ATP]_mito_ **(B)** and [Ca^2+^]_i_ **(C)** transients in female (magenta) and male (blue) myocytes. Data are presented as mean ± SD (female: N = 3 mice, *n* = 26 and 17 cells of Mode 1 and 2 sites, respectively; male: N = 3 mice, *n* = 25 and 24 cells of Mode 1 and 2 sites, respectively). All significant values are provided from a nested *t* test. Light symbols: individual cells; dark symbols: individual animals.

To assess sex-dependent differences in the kinetics of mitochondrial ATP transients, we quantified time-to-peak and time-to-decay for Mode 1 and Mode 2 responses (**SI Figure 1A, B**). In Mode 1 sites, [ATP]_mito_ rose faster in females (80 ± 23 ms) than in males (97 ± 25 ms; *P* = 0.0453), while Mode 2 rise times were similar between sexes (females: 107 ± 43 ms; males: 106 ± 45 ms; P = 0.9662) (**SI Figure 1C**). Time to 90% decay was also comparable across sex and mode (**SI Figure 1D**) (Mode 1: 123 ± 51 and 135 ± 56 ms in female and male, respectively, P = 0.4820; Mode 2: 144 ± 65 and 121 ± 60 ms in female and male, respectively, P = 0.4025). The time-to-peak and time-to-decay values for [Ca^2+^]_i_ transients were similar in Mode 1 and Mode 2 sites across both sexes (**SI Figure 1E, F**) (Time to peak: Mode 1 - 59 ± 9 and 63 ± 14 ms in female and male, respectively, P = 0.2201; Mode 2 - 61 ± 12 and 60 ± 11 ms in female and male, respectively, P = 0.8721) (Time to decay: Mode 1 - 228 ± 70 and 226 ± 50 ms in female and male, respectively, P = 0.9676; Mode 2 - 233 ± 62 and 209 ± 43 ms in female and male, respectively, P = 0.4244). It is important to note that, while [Ca^2+^]_i_ and [ATP]_mito_ transients are shown for temporal alignment, a direct comparison of their kinetics should be interpreted with caution, given the markedly faster response properties of Rhod-3 compared with those of the EGFP-based mito-iATP sensor.

To exclude motion artifacts and validate the pseudo-ratiometric approach, we labeled the HaloTag domain of mito-iATP with the red-shifted JFX554 HaloTag ligand and simultaneously imaged both channels during field stimulation. Robust beat-locked Mode 1 and Mode 2 [ATP]_mito_ transients were evident in the mito-iATP channel, whereas the JFX554 HaloTag signal exhibited no detectable beat-synchronous changes in line-scan recordings (**Figure 4A, C**). Across regions exhibiting ATP gains or dips, the mito-iATP ratio signal scaled linearly with the underlying non-ratiometric mito-iATP fluorescence (**Figure 4B, D**), indicating that the ratio faithfully reports changes in matrix ATP rather than movement of mitochondria within the confocal volume. These data argue against motion or volume changes as the source of the observed [ATP]_mito_ waveforms and support the use of mito-iATP ratios to quantify beat-locked microdomain ATP dynamics.

**Figure 4.**
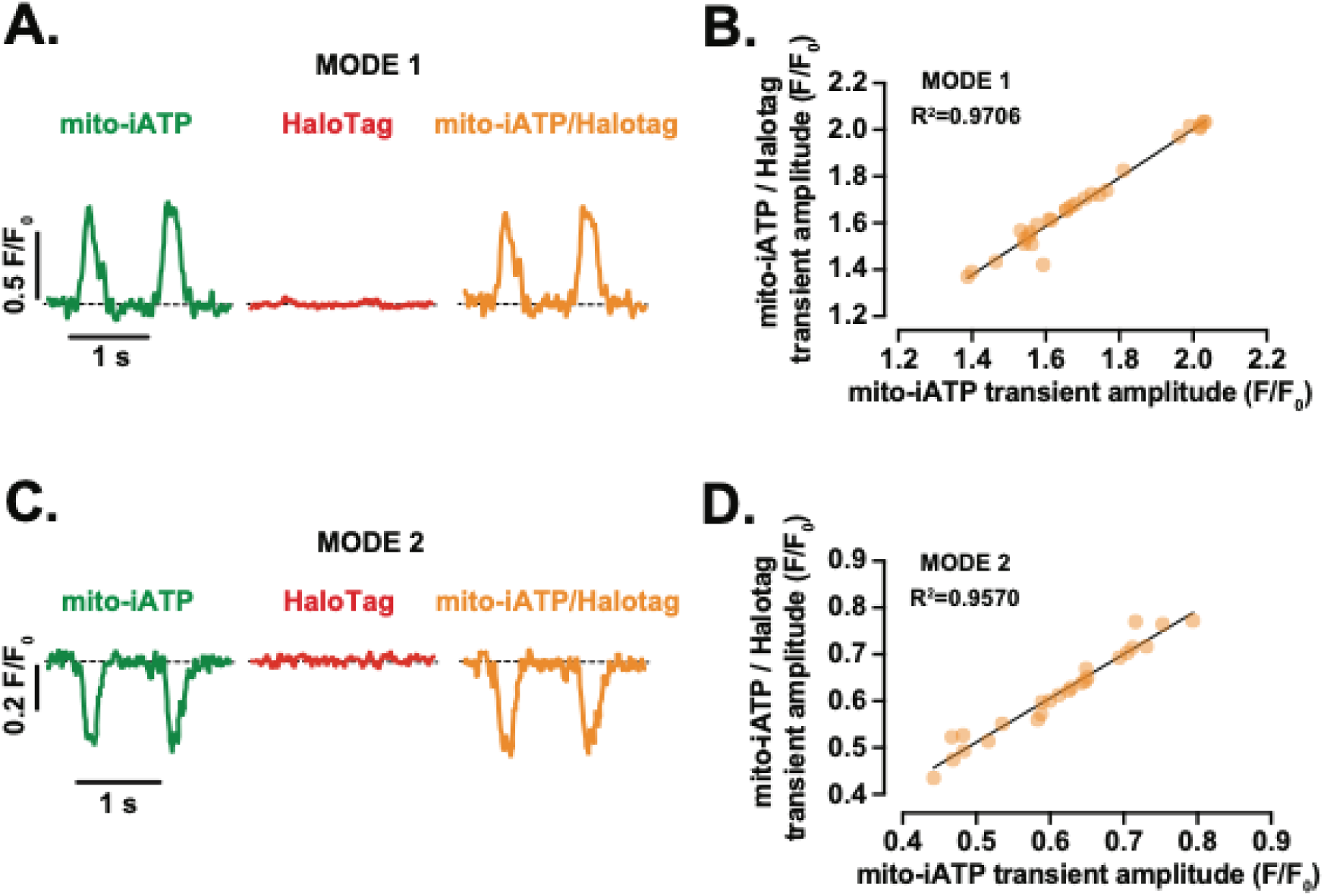
Ratiometric validation confirms that beat-to-beat mitochondrial iATP transients are independent of motion artifacts. **(A,C)** Representative time course from line scans of [ATP]_mito_ (green), reference fluorophore JFX554 HaloTag (red), and ratiometric signal (orange) from myocytes with Mode 1 **(A)** and Mode 2 **(C)** ATP dynamics. All signals are shown as normalized fluorescence (F/F_0_). **(B,D)** Relationship between [ATP]_mito_ transient amplitude and ratiometric transient amplitude for Mode 1 **(B)** and Mode 2 **(D)** events. Points indicate individual transients, and line shows the fitted relationship with the coefficient of determination (R^2^) reported on-graph. Data are presented as mean ± SD (female: N = 2 mice n = 6 cells/14 transients and 5 cells/18 transients of Mode 1 and 2 sites, respectively; male: N = 2 mice, *n* = 4 cells/14 transients and 3 cells/7 transients of Mode 1 and 2 sites, respectively).

Together, these findings demonstrate that millisecond-scale ATP transients can be visualized within adult ventricular mitochondria and that they adopt two discrete, spatially confined waveforms. While peak amplitudes are similar, the relative distribution, spatial heterogeneity, and kinetics of these events differ between male and female myocytes, raising the possibility that sex-specific energetic strategies reflect both mitochondrial abundance and subcellular organization.

### MCU and ANT are required for beat-locked mitochondrial ATP transients

To determine whether mitochondrial Ca^2+^ uptake through the mitochondrial Ca^2+^ uniporter (MCU) is required for the beat-locked [ATP]_mito_ signals, ventricular myocytes expressing mito-iATP and loaded with Rhod-3 were exposed to 10 µM Ru360, a selective MCU blocker (**Figure 5A, B, E, F**). Under control conditions, each action potential evoked robust Mode 1 or Mode 2 [ATP]_mito_ transients (Mode 1: 1.60 ± 0.13 and 1.59 ± 0.21 F/F_0_; Mode 2: 0.68 ± 0.05 and 0.69 ± 0.05 F/F_0_; female and male, respectively) that were tightly phase-locked to the [Ca^2+^]_i_ transient (Mode 1: 3.03 ± 0.69 and 2.73 ± 0.69 F/F_0_; Mode 2: 2.96 ± 0.53 and 2.65 ± 0.48 F/F_0_; female and male, respectively), as described above. Exposure to Ru360 decresed these beat-coupled [ATP]_mito_ oscillations in both modes, markedly reducing local ATP fluctuations to near-baseline noise (**Figure 5C, G**) (Mode 1: 1.22 ± 0.18 F/F_0_ in female, P = 0.0013 vs control; and 1.24 ± 0.11 F/F_0_ in males, P = 0.0031 vs control; Mode 2: 0.79 ± 0.05 F/F_0_ in female, P = 0.0082 vs control; and 0.78 ± 0.05 F/F_0_ in males, P = 0.0122 vs control), while simultaneously recorded [Ca^2+^]_i_ transients were maintained (**Figure 5D, H**) (Mode 1-associated Ca^2+^ signal: 2.06 ± 0.32 F/F_0_ in female, P = 0.0041 vs control; and 2.04 ± 0.57 F/F_0_ in males, P = 0.0381 vs control; Mode 2-associated Ca^2+^ signal: 1.97 ± 0.46 F/F_0_ in female, P = 0.0093 vs control; and 2.07 ± 0.53 F/F_0_ in males, P = 0.0459 vs control). Thus, MCU-mediated Ca^2+^ entry is critical for generating the phasic mitochondrial ATP responses during EC coupling.

**Figure 5.**
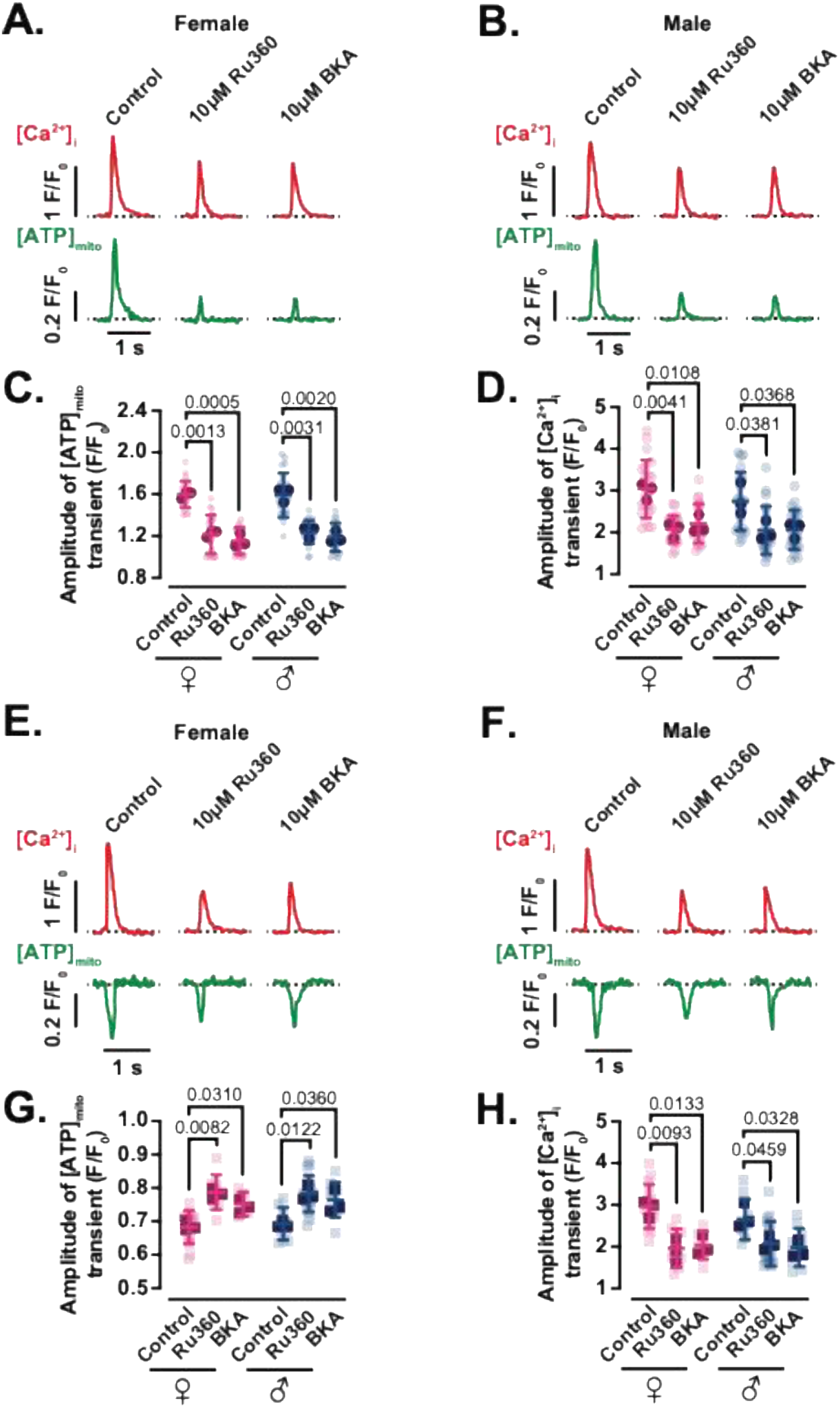
Beat-to-beat metabolic signaling requires mitochondrial Ca^2+^ uptake and ANT-mediated ATP export. Representative time course from line scans of [Ca^2+^]_i_ (top) and [ATP]_mito_ (bottom) in female **(A)** and male **(B)** myocytes with Mode 1 ATP dynamics in control cells and in cells exposed to 10 μM of BKA or Ru360 for 30 minutes. Scatter plots of the amplitudes of [ATP]_mito_ **(C)** and [Ca^2+^]_i_ **(D)** transients in female (magenta) and male (blue) myocytes with Mode 1 ATP dynamics under control and treated conditions (female: N = 3 mice, *n* = 22, 20 and 22 cells; male: N = 3 mice, *n* = 21, 23 and 22 cells. Data from control, BKA and Ru360 conditions, respectively). Representative time course from line scans of [Ca^2+^]_i_ (top) and [ATP]_mito_ (bottom) in female **(E)** and male **(F)** myocytes with Mode 2 ATP dynamics in control cells and in cells exposed to 10 μM of BKA or Ru360 for 30 minutes. Scatter plots of the amplitudes of [ATP]_mito_ **(G)** and [Ca^2+^]_i_ **(H)** transients in female (magenta) and male (blue) myocytes with Mode 2 ATP dynamics under control and treated conditions (female: N = 3 mice, *n* = 15, 7 and 9 cells; male: N = 3 mice, *n* = 10, 10 and 18 cells. Data from control, BKA and Ru360 conditions, respectively). Data are presented as mean ± SD. All significant values are provided from a nested *t* test. Light symbols: individual cells; dark symbols: individual animals.

We next tested whether export of matrix ATP via the adenine nucleotide translocase (ANT) is necessary to sustain beat-locked ATP signals in the mitochondrial matrix (**Figure 5A, B, E, F**). Inhibition of ANT with bongkrekic acid (BKA; 10 µM) reduced the beat-synchronized [ATP]_mito_ transients (**Figure 5C, G**) (Mode 1: 1.16 ± 0.13 F/F_0_ in female, P = 0.0005 vs control; and 1.19 ± 0.13 F/F_0_ in males, P = 0.0020 vs control; Mode 2: 0.75 ± 0.04 F/F_0_ in female, P = 0.0310 vs control; and 0.76 ± 0.05 F/F_0_ in males, P = 0.0360 vs control) maintaining [Ca^2+^]_i_ transients (**Figure 5D, H**) (Mode 1: 2.20 ± 0.46 F/F_0_ in female, P = 0.0108 vs control; and 2.05 ± 0.47 F/F_0_ in males, P = 0.0368 vs control; Mode 2: 2.04 ± 0.34 F/F_0_ in female, P = 0.0133 vs control; and 1.99 ± 0.45 F/F_0_ in males, P = 0.0328 vs control), indicating that continued nucleotide exchange across the inner mitochondrial membrane is required for the mito-iATP signal. Together, these interventions demonstrate that beat-to-beat ATP transients in mitochondrial matrix require MCU-dependent mitochondrial Ca^2+^ uptake to drive oxidative phosphorylation and ANT-mediated nucleotide exchange to transmit this oscillatory ATP output to the cytosol.

### Female myocytes trade mitochondrial mass for tighter SR–mitochondrial coupling and higher Mfn2/ATP5A density

We postulated that the smaller local ATP signals and lower prevalence of ATP-gain microdomains observed in male myocytes might stem from sex differences in both mitochondrial content and the spatial relationship between mitochondria and sites of SR Ca^2+^ release. Voxel segmentation of 3D confocal stacks from Mitotracker-labeled ventricular myocytes revealed that mitochondria occupied 6391 ± 1700 μm^3^ of cytosolic volume in females versus 8192 ± 1264 μm^3^ in males (n = 30 cells per sex; P = 0.0307), indicating that male myocytes devote a larger fraction of their cytosol to mitochondrial volume (**Figure 6A, B**).

**Figure 6.**
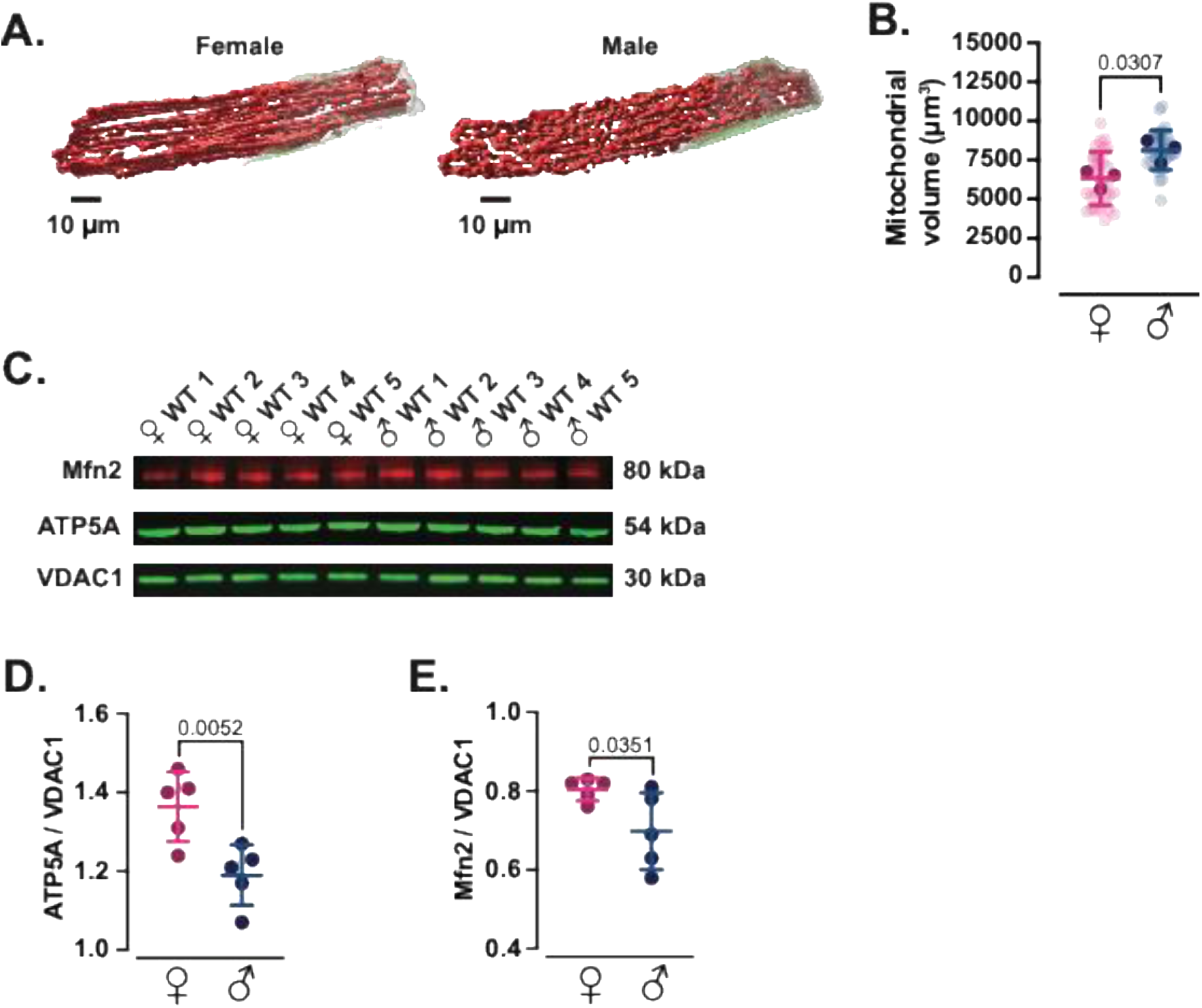
Female myocytes harbor a smaller mitochondrial volume but exhibit tighter SR–mitochondrial coupling. **(A)** 3D reconstructions of the mitochondrial volume in a female (left) and male (right) ventricular myocyte. Mitochondria were volumetrically rendered from deconvolved confocal z-stacks after staining with MitoTracker Deep Red. **(B)** Scatter plot of the mitochondrial volume in female (magenta) and male (blue) myocytes (female and male: N = 3 mice, 30 cells). Significant values are provided from a nested *t* test. **(C)** Representative immunoblots of the mitochondrial outer-membrane tether protein mitofusin 2 (Mfn2, 80 kDa), of the α subunit of the catalytic core of mitochondrial ATP synthase protein (ATP5A, 54 kDa) and of the voltage-dependent anion channel 1 (VDAC1, 30kDa) in ventricular lysates from five female (♀ WT 1–5) and five male (♂ WT 1–5) mice. Densitometric quantification of ATP5A **(D)** and Mfn2 **(E)**, normalized to VDAC1. Significant values are provided from a Student’s *t* test. All data are presented as mean ± SD. Light symbols: individual cells; dark symbols: individual animals.

To explore the molecular correlates of this architectural divergence, we quantified protein levels of mitofusin-2 (Mfn2), a key tether that links mitochondria to the SR and facilitates Ca^2+^-dependent stimulation of oxidative phosphorylation (Chen *et al*., 2012; Dorn *et al*., 2015; Rhana *et al*., 2024), and ATP5A, a core subunit of the mitochondrial ATP synthase complex. Immunoblotting of ventricular homogenates showed that levels of ATP5A (1.36 ± 0.09 in female, 1.19 ± 0.08 in male; P = 0.0052) and Mfn2 (0.80 ± 0.03 in female, 0.70 ± 0.10 in male; P = 0.0351) were higher in female than male when normalized per mitochondrial density (**Figure 6C-E**).

We next asked whether this inferred molecular enrichment is accompanied by tighter physical coupling between mitochondria and SR Ca^2+^ release sites (**Figure 7**). Ventricular myocytes were co-immunolabeled with antibodies against the outer mitochondrial membrane protein VDAC and the SR Ca²⁺ pump SERCA, and confocal images from each channel were independently thresholded to generate binary masks (**Figure 7A**). These binary images were then multiplied pixel-by-pixel so that only voxels positive for both VDAC and SERCA were set to 1; in the resulting overlap maps, colocalized regions appear as discrete black puncta, whose total area was quantified as an index of SR–mitochondrial apposition. This analysis revealed a significantly larger VDAC–SERCA overlap area in female than male myocytes (**Figure 7-B**) (59 ± 7 % in female vs 47 ± 9 % in male, P = 0.0324), indicating that a greater fraction of mitochondrial surface resides in close proximity to SR Ca^2+^ handling machinery in females.

**Figure 7.**
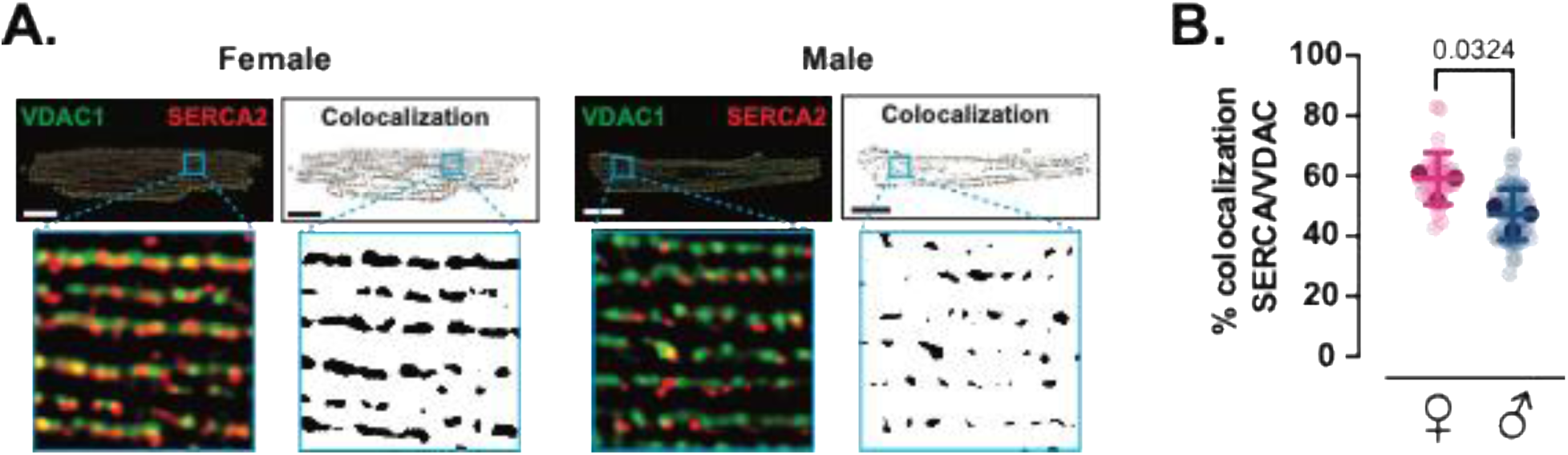
SERCA2-VDAC1 colocalization reveals sex-dependent SR–mitochondria proximity. **(A)** Representative confocal immunofluorescence images from female and male ventricular myocytes labeled for the mitochondrial outer membrane marker VDAC1 (green) and the sarcoplasmic reticulum Ca²⁺-ATPase SERCA2 (red). Merged images and 10μm x 10μm insets are shown alongside a binary colocalization maps that shows pixels in which SERCA2 and VDAC1 are completely overlapped. Scale bars: 10μm. **(B)** Scatter plot of SERCA2-VDAC1 percent colocalization in female (magenta) and male (blue) myocytes. Data are presented as mean ± SD (female: N = 3 mice, 33 cells; male: N = 3 mice, 51 cells). All significant values are provided from a nested *t* test. Light symbols: individual cells; dark symbols: individual animals.

Together with the mito-iATP imaging, these data support a model in which female myocytes trade mitochondrial mass for more densely equipped and more tightly SR-coupled mitochondria, favoring rapid, spatially confined ATP surges in response to local Ca^2+^ fluxes, whereas male myocytes rely more heavily on increased mitochondrial volume to scale ATP output. This structural and molecular arrangement dovetails with our prior finding that beat-locked increases in mitochondrial Ca^2+^ are tightly associated with cytosolic iATP transients in ventricular myocytes (Rhana *et al*., 2024), suggesting that enhanced SR–mitochondrial tethering and higher Mfn2/ATP5A density in females sharpen this Ca^2+^-to-ATP coupling axis.

### Cytosolic ATP transients mirror mitochondrial waveforms and exhibit a sex-dependent domain organization

To assess cytosolic ATP dynamics, we used iATPSnFR1, the genetically encoded cytosolic fluorescent ATP sensor developed by Lobas *et al*. (2019) (**Figure 8**). The construct, here termed cyto-iATP, was packaged into an AAV9 vector and expressed in adult ventricular myocytes from male and female mice. Confocal imaging showed a uniform, diffuse fluorescence pattern throughout the cytosol in male or female myocytes (**Figure 8A, B**).

**Figure 8.**
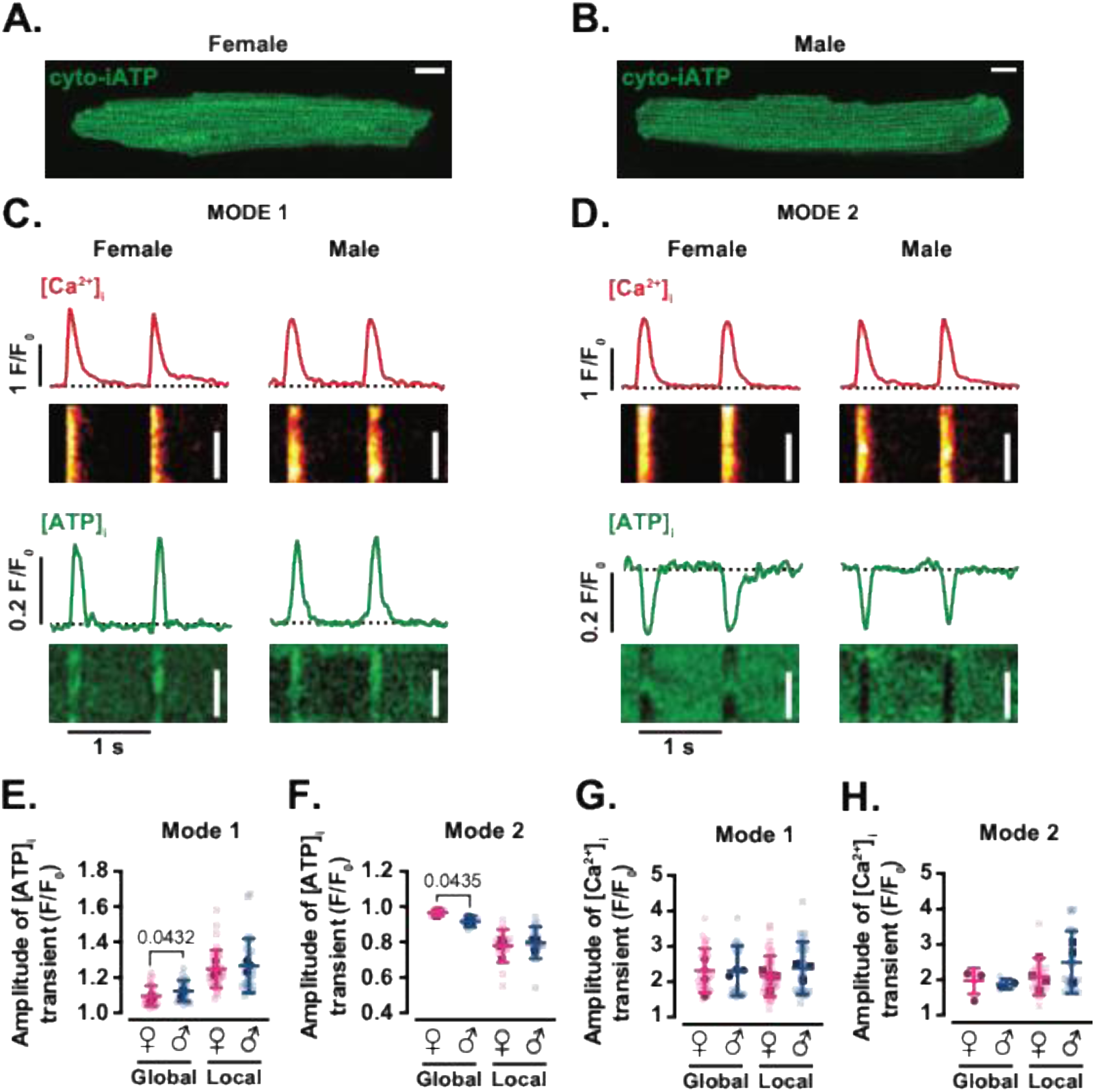
Cytosolic ATP transients mirror mitochondrial modes and show sex-dependent domain organization. Confocal image of an exemplar ventricular myocyte expressing cytosolic iATP sensor in female **(A)** and male **(B)** mice. Scale bar: 10 μm. Representative line scan images of [Ca^2+^]_i_ (top) and [ATP]_i_ (bottom) in female and male myocytes with Mode 1 **(C)** or Mode 2 **(D)** ATP dynamics. Red and green traces above each line-scan show the time course of [Ca^2+^]_i_ and [ATP]_i_ transients, respectively. Scale bars: 20 µm. Scatter plots of global and local amplitudes of [ATP]_i_ transients in Mode 1 **(E)** and Mode 2 **(F)** sites in female (magenta) and male (blue) myocytes. Scatter plots of global and local amplitudes of [Ca^2+^]_i_ transients in Mode 1 **(G)** and Mode 2 **(H)** sites in female (magenta) and male (blue) myocytes. Data are presented as mean ± SD (female: N = 4 mice, n = 24 cells/35 sites and 4 cells/16 sites of Mode 1 and 2 (global/local), respectively; male: N = 3 mice, n = 20 cells/26 sites and 5 cells/17 sites of Mode 1 and 2 (global/local), respectively). All significant values are provided from a nested *t* test. Light symbols: individual cells; dark symbols: individual animals.

To determine whether [ATP]_i_ exhibits beat-to-beat fluctuations during EC coupling, we recorded simultaneous line-scan images of cyto-iATP and Rhod-3 fluorescence in cells subject to field stimulation (1 Hz). As with mitochondrial ATP, [ATP]_i_ underwent rhythmic phasic changes that were tightly phase-locked to the [Ca^2+^]_i_ transient. Two distinct waveform types—Mode 1 and Mode 2—were identified. In Mode 1 (**Figure 8C**), [ATP]_i_ rose rapidly after the Ca^2+^ transient and returned to baseline before the next beat. In Mode 2 (**Figure 8D**), [ATP]_i_ dipped transiently and recovered with similar timing. Both modes were present within single cells and segregated spatially, revealing a cytosolic energetic microdomain structure analogous to that observed in the mitochondrial matrix.

Quantification of local and global [ATP]_i_ and [Ca^2+^]_i_ transients revealed striking parallels to the mitochondrial data (**Figure 8E–H**). Mode 1 and Mode 2 events were detected in myocytes of both sexes, and their spatial distribution followed a similar sex-dependent pattern: female myocytes exhibited a higher proportion of ATP-gain (Mode 1) domains (58% in females and 40% in males), whereas male myocytes had more ATP-dip (Mode 2) regions (10% in females and 13% in males). Local transient amplitudes did not differ significantly between sexes for either mode (**Figure 8E, F**) (Mode 1: 1.25 ± 0.11 F/F_0_ in female and 1.27 ± 0.15 F/F_0_ in male, P = 0.6054; Mode 2: 0.77 ± 0.09 F/F_0_ in female and 0.79 ± 0.09 F/F_0_ in male, P = 0.5789), indicating that the intrinsic dynamics of each waveform are comparable. However, global F/F_0_ signals were significantly larger in male than female myocytes for both Mode 1 (1.09 ± 0.05 vs 1.13 ± 0.06 F/F_0_; *P* = 0.0432) and Mode 2 (0.96 ± 0.02 vs 0.91 ± 0.03 F/F_0_; *P* = 0.0435), consistent with a greater fraction of the cytosol participating in each mode in male cells. Thus, the amplitude of [Ca^2+^]_i_ transients was similar in cells displaying either Mode 1 (Global: 2.23 ± 0.57 F/F_0_ in female and 2.30 ± 0.72 F/F_0_ in male, *P* = 0.7489; Local: 2.15 ± 0.58 F/F_0_ in female and 2.38 ± 0.75 F/F_0_ in male, *P* = 0.2335) or Mode 2 fluctuations (**Figure 8G, H**) (Global: 1.96 ± 0.37 F/F_0_ in female and 2.17 ± 0.59 F/F_0_ in male, *P* = 0.5469; Local: 2.08 ± 0.53 F/F_0_ in female and 2.49 ± 0.88 F/F_0_ in male, *P* = 0.2746) across both sexes.

To evaluate sex- and mode-dependent differences in the kinetics of [ATP]_i_ transients, we measured the time to peak and decay in Mode 1 and Mode 2 regions of male and female myocytes (**SI Figure 2**). Unlike [ATP]_mito_ transients, [ATP]_i_ transients in Mode 1 sites showed no significant difference in the time to 90% peak between sexes (87 ms for both, P = 0.9962). However, time to 90% peak [ATP]_i_ at Mode 2 sites differed between female (75 ± 20 ms) and male (107 ± 48 ms) myocytes (*P* = 0.0448). As was the case for [ATP]_mito_ transients, we observed no difference in time-to-decay of [ATP]_i_ transients across sexes and modes or the kinetics of whole-cell [Ca^2+^]_i_ transients.

### Cytosolic ATP transients persist under a mixed oxidative substrate supply

To test whether beat-locked cytosolic ATP dynamics are preserved under more physiological oxidative substrates, we recorded [Ca^2+^]_i_ and [ATP]_i_ in field-stimulated ventricular myocytes (1 Hz) perfused with a substrate mix containing pyruvate, lactate, carnitine, and palmitate. Representative line-scan time courses show that Mode 1 cells maintained beat-synchronous [ATP]_i_ increases aligned with [Ca^2+^]_i_ transients, whereas Mode 2 cells retained beat-locked [ATP]_i_ decreases (ATP “dips”) despite robust [Ca^2+^]_i_ transients, in both female and male myocytes (**Figure 9A**). Like with mito-iATP, the amplitude of [ATP]_i_ transients in both Mode 1 and 2 ATP dynamics were indistinguishable from those recorded in Tyrode’s solution (**Figure 9B**) (Mode 1: 1.30 ± 0.13 F/F_0_ in female, P = 0.1649 vs control; and 1.30 ± 0.12 F/F_0_ in male, P = 0.5957 vs control; Mode 2: 0.76 ± 0.10 F/F_0_ in female, P = 0.7108; and 0.79 ± 0.08 F/F_0_ in male, P = 0.9373 vs control).

**Figure 9.**
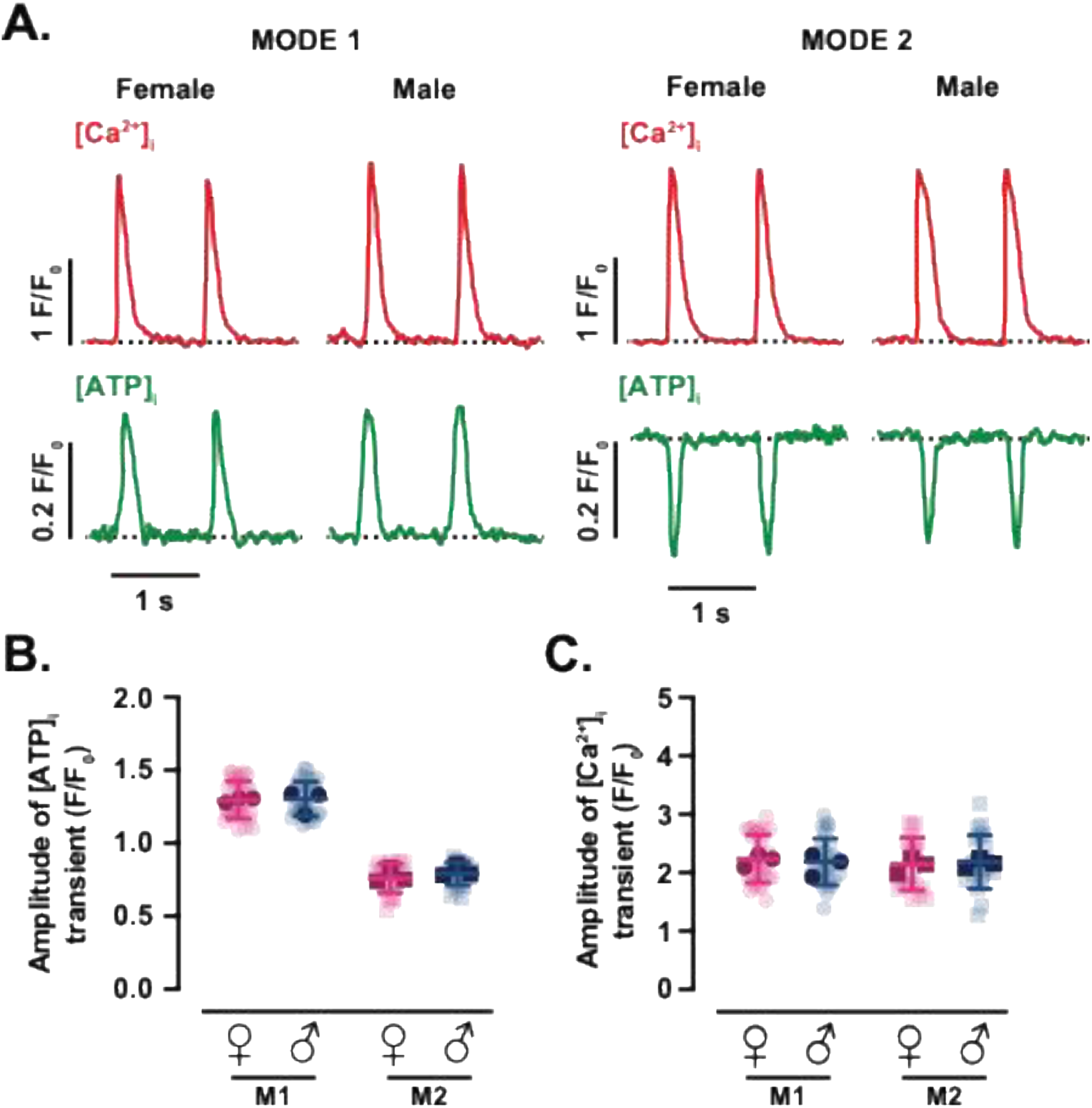
Effect of mixed substrates on cytosolic ATP and [Ca²⁺]_i_ transient measurements. **(A)** Representative time course from line scans of [Ca^2+^]_i_ (top) and [ATP]_i_ (bottom) in female and male myocytes with Mode 1 (left) or Mode 2 (right) ATP dynamics using a new physiological substrate mix containing pyruvate, lactate, carnitine, and palmitate. Scatter plots of the amplitudes of [ATP]_i_ **(B)** and [Ca^2+^]_i_ **(C)** transients in female (magenta) and male (blue) myocytes. Data are presented as mean ± SD (female: N = 3 mice, *n* = 30 and 21 cells of Mode 1 and 2 sites, respectively; male: N = 3 mice, *n* = 27 and 22 cells of Mode 1 and 2 sites, respectively). All significant values are provided from a nested *t* test. Light symbols: individual cells; dark symbols: individual animals.

In parallel, [Ca^2+^]_i_ transient amplitudes were comparable across sexes and modes (**Figure 9C**) (Mode 1: 2.24 ± 0.41 F/F_0_ in female, P = 0.5033 vs control; and 2.19 ± 0.40 F/F_0_ in male, P = 0.3075 vs control; Mode 2: 2.15 ± 0.45 F/F_0_ in female, P = 0.6893; and 2.18 ± 0.46 F/F_0_ in male, P = 0.3474 vs control), indicating that the preservation of ATP transients in this substrate condition is not attributable to gross differences in [Ca^2+^]_i_ transient magnitude. These data show that cytosolic ATP transients are preserved under pyruvate/lactate/carnitine/palmitate and are qualitatively similar to those observed in Tyrode solution, supporting the view that Mode 1 and 2 ATP dynamics exist across substrate environments.

### Cross-bridge inhibition selectively attenuates Mode 2 cytosolic ATP dips while preserving Mode 1 gains

Because cross-bridge cycling is a major ATP sink during each beat, we next asked whether contractile activity contributes to the Mode 2 cytosolic ATP phenotype. We therefore applied 10 µM blebbistatin to inhibit myosin II ATPase and uncouple shortening from the Ca^2+^ transient (**Figure 10**). Blebbistatin decreased contraction by 74 ± 16% (P < 0.0001). Representative line-scan recordings show that beat-locked [ATP]_i_ transients in both female and male myocytes exhibiting Mode 1 gains or Mode 2 dips even in the presence of blebbistatin (**Figure 10A, B**).

**Figure 10.**
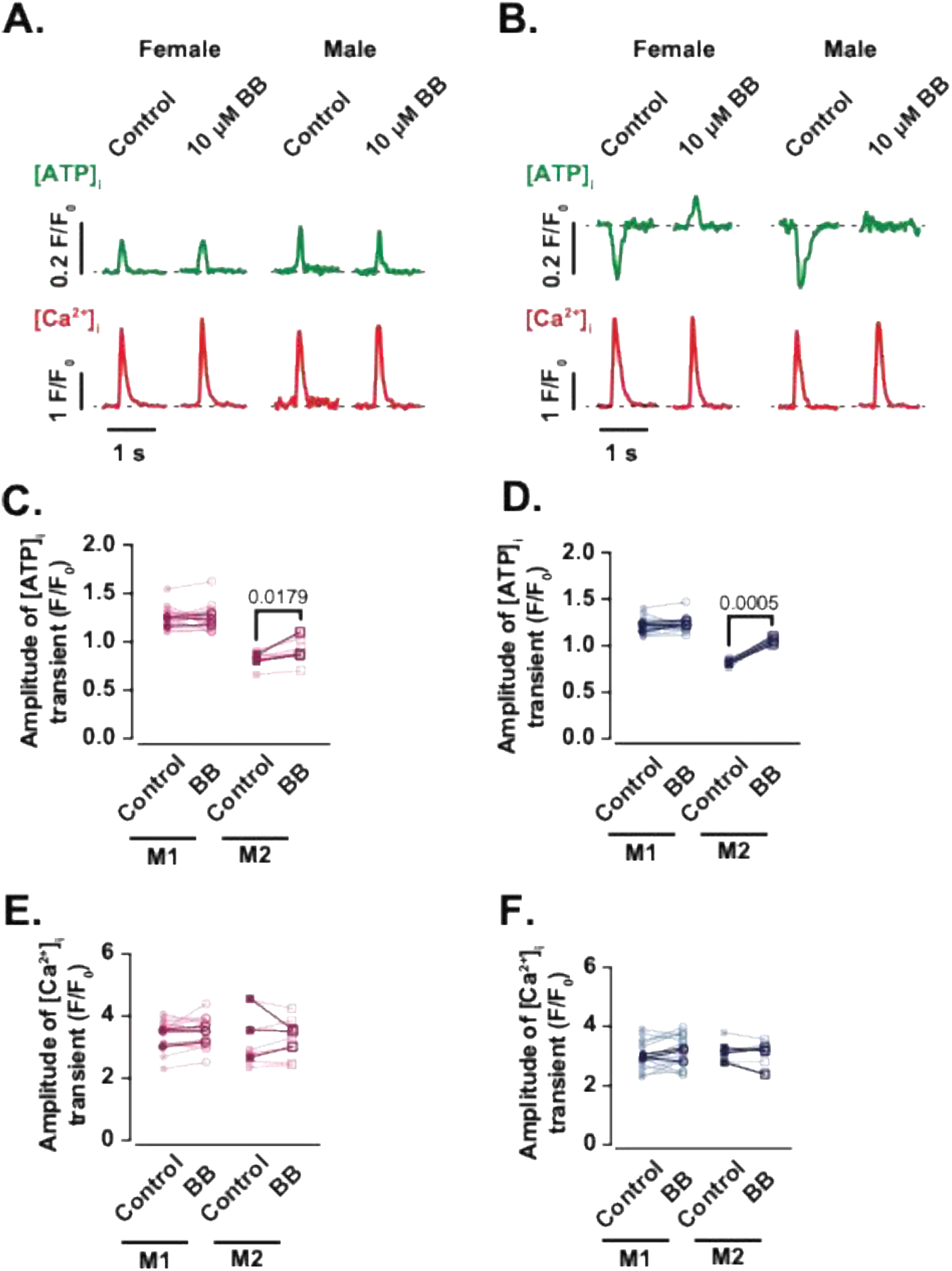
Blebbistatin suppresses contraction while preserving beat-to-beat cytosolic ATP and [Ca²⁺]_i_ transients. Representative time course from line scans of [ATP]_i_ (green) and [Ca^2+^]_i_ (red) in female and male myocytes with Mode 1 **(A)** or Mode 2 **(B)** ATP dynamics in control condition and in the presence of 10 μM Blebbistatin (BB) for 30 minutes. Scatter plots of the amplitude of [ATP]_i_ **(C, D)** and [Ca^2+^]_i_ **(E, F)** transients of Mode 1 and 2 ATP dynamics under control and treated conditions in female (magenta) and male (blue) myocytes. Data are presented as mean ± SD (female: N = 3 mice, *n* = 13 and 6 cells of Mode 1 and 2 sites, respectively; male: N = 3 mice, *n* = 12 and 5 cells of Mode 1 and 2 sites, respectively). All significant values are provided from a paired *t* test. Light symbols: individual cells; dark symbols: individual animals.

Notably, while the amplitude of Mode 1 [ATP]_i_ transients was largely preserved (1.26 ± 0.12 vs 1.24 ± 0.14 F/F_0_ in female, P = 0.3276; and 1.22 ± 0.09 vs 1.23 ± 0.09 F/F_0_ in male, P = 0.2310), Mode 2 [ATP]_i_ dips were significantly attenuated in both females (0.83 ± 0.09 vs 0.92 ± 0.14 F/F_0_, P = 0.0179) and males (0.82 ± 0.04 vs 1.05 ± 0.04 F/F_0_, P = 0.0005) (**Figure 10C, D)**. The [Ca^2+^]_i_transient amplitude showed no evident change with blebbistatin in either mode or sex (**Figure 10E, F**). These results, consistent with our prior observations in ventricular myocytes (Rhana *et al*., 2024), support the view that Mode 2 reflects local ATP depletion driven by contractile ATP utilization, whereas Mode 1 gains persist when cross-bridge ATP demand is reduced.

### Increasing workload elevates [ATP]_i_ more rapidly and to a greater extent in male than female myocytes

To determine whether the ∼20% lower mitochondrial volume in female ventricular myocytes slows cytosolic ATP replenishment during increased workload, we monitored diastolic [ATP]_i_ in cells paced at varying frequencies (**Figure 11**). Upon switching from quiescence (0 Hz) to 2 Hz stimulation, diastolic [ATP]_i_ rose and stabilized at a higher set point that persisted throughout pacing (**Figure 11A–C**). The new diastolic [ATP]_i_ values (in F/F_0_ units) were 1.22 ± 0.11 in female and 1.25 ± 0.07 in male myocytes (P = 0.8967) (**Figure 11D**). Notably, further increasing pacing to 3 Hz produced no additional rise (P = 0.9644), suggesting a physiological ceiling to workload-induced ATP enhancement.

**Figure 11.**
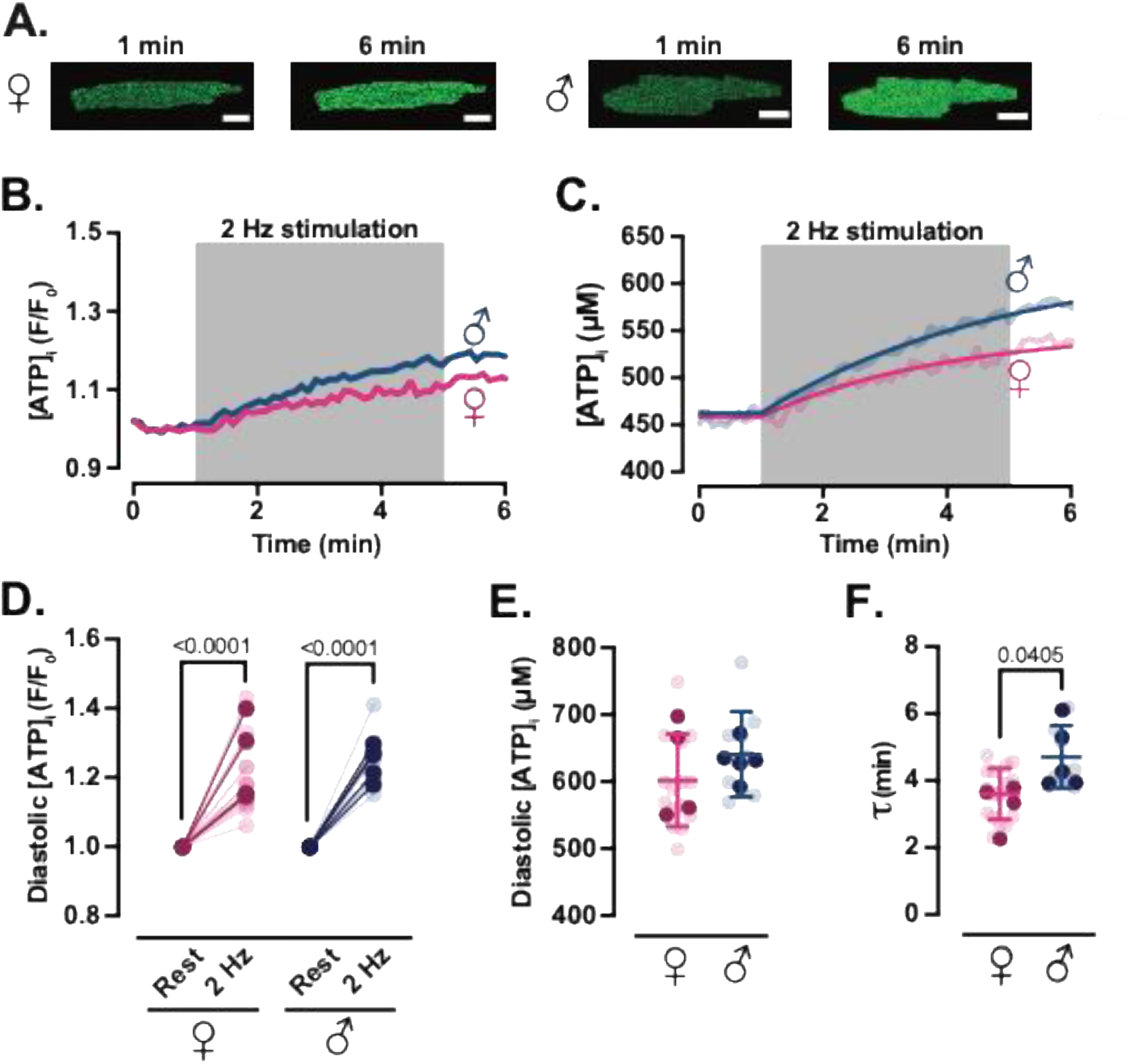
Diastolic cytosolic ATP rises with increased pacing frequency. **(A)** Representative confocal images of female (♀, left) and male (♂, right) ventricular myocytes expressing cyto-iATP. Fluorescence is shown before (1 minute) and after (6 minutes) 2 Hz field stimulation. Scale bars, 10 µm. Representative time course of [ATP]_i_ in normalized fluorescence (F/F₀) **(B)** and in μM units **(C)** during the 6-minute experiment in female (magenta) and (male) myocytes. The gray box indicates the 2-Hz stimulation period (1–5 minutes). F/F_0_ data were converted to intracellular ATP concentration (µM) using a pseudo-ratiometric method. **(D)** Scatter plot of diastolic [ATP]_i_ before and after 2 Hz stimulation as normalized fluorescence (F/F_0_). **(E)** Scatter plot of diastolic [ATP]_i_ at the end of 2 Hz stimulation as concentration (µM). **(F)** Scatter plot of the time constant for the rise in [ATP]_i_ during pacing. Data are expressed as means ± SD (female: N = 4 mice, *n* = 16 cells; male: N = 5 mice, *n* = 10 cells). All significant values are provided from a nested *t* test. Light symbols: individual cells; dark symbols: individual animals.

Because the relationship between absolute cytosolic [ATP]_i_ and cyto-iATP fluorescence is nonlinear (Lobas *et al*., 2019; Rhana *et al*., 2024), we converted F/F_0_ values to estimated ATP concentrations using experimentally determined diastolic cytosolic [ATP]. We measured diastolic [ATP]_i_ in female myocytes (471 ± 81 µM) and found it to be similar to that in male myocytes (457 ± 199 µM; P = 0.6169) (Rhana *et al*., 2024). Although the 2 Hz-evoked mean diastolic [ATP]_i_ trended lower in female (603 ± 69 μM) than male (642 ± 63 μM) myocytes, this difference was not statistically significant (P =0.5303), indicating comparable resting ATP levels between sexes (**Figure 11E**).

We then fit the time course of the diastolic [ATP]_i_ rise (in μM) using a single-exponential function. Kinetic fits were applied to the estimated [ATP]_i_ rather than F/F_0_ to avoid nonlinearity-induced distortions in estimating time constants. This analysis revealed that female myocytes reached their new steady-state ATP level with a time constant (τ) of 3.62 ± 0.77 minutes, whereas male myocytes required ∼23% longer (τ = 4.72 ± 0.94 minutes; *P* = 0.0405) (**Figure 11F**). This sex difference supports the hypothesis that tighter SR–mitochondrial coupling in female myocytes enables more rapid activation of oxidative phosphorylation, while weaker coupling in male myocytes introduces a delay.

Together, these data suggest that the diastolic component of the cytosolic ATP response to increased workload is governed by intrinsic kinetic programs modulated by both mitochondrial reserve and SR–mitochondrial coupling. Female myocytes achieve faster ATP upregulation despite reduced mitochondrial content, whereas male myocytes rely on greater mitochondrial mass but exhibit slower metabolic adaptation.

### Differential modulation of beat-to-beat [ATP]_i_ transients by pacing frequency in male and female myocytes

To test how acute workload modulates cytosolic ATP oscillations, we imaged cyto-iATP signals in ventricular myocytes paced first at 1 Hz and then at 2 Hz (**Figure 12A, B**). We selected 2 Hz because diastolic [ATP]_i_ rises in response to this—but not higher—frequencies, as we mentioned in **Figure 11**, making it the highest rate that reliably elevates energetic demand while preserving cellular integrity. Because diastolic [ATP]_i_ is higher at 2 Hz, beat-evoked F/F_0_ values were converted to concentration units using sex-specific baselines (male, 642 µM; female, 603 µM).

**Figure 12.**
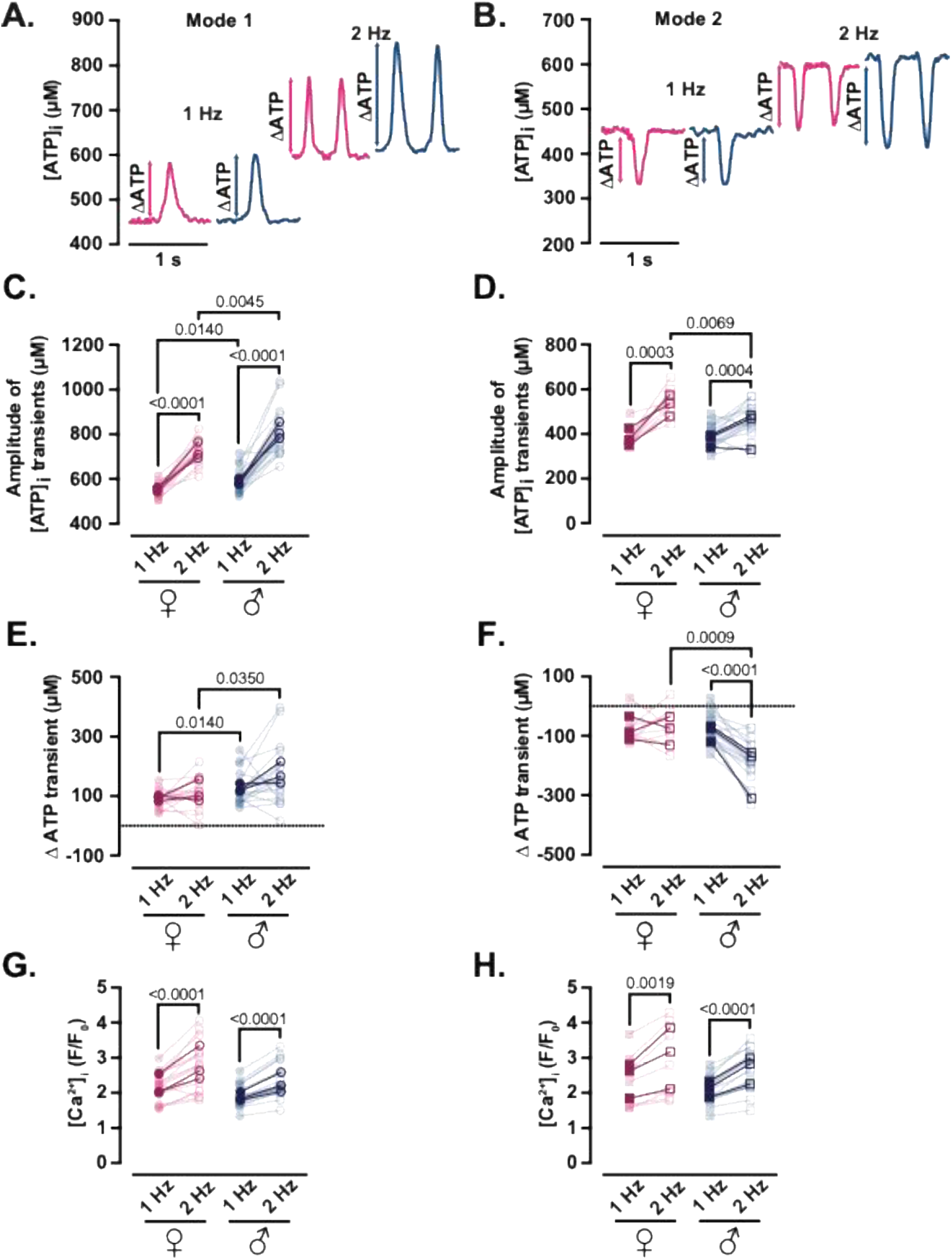
Increased workload amplifies cytosolic ATP transients in male but not female myocytes. Representative time course of [ATP]_i_ from line scans of female (magenta) and male (blue) myocytes with Mode 1 **(A)** and Mode 2 **(B)** ATP dynamics field-stimulated at 1 Hz (left) or 2 Hz (right). Double-headed arrows indicate the net shift in [ATP]_i_ consumption or production (ΔATP) after increasing workload (2 Hz). Scatter plots of the amplitudes of [ATP]_i_ transients of female (magenta) and male (blue) myocytes with Mode 1 **(C)** and Mode 2 **(D)** ATP dynamics under 1-2Hz stimulation. Scatter plots of ΔATP transients of female (magenta) and male (blue) myocytes with Mode 1 **(E)** and Mode 2 **(F)** ATP dynamics under 1-2 Hz stimulation. Scatter plots of the amplitudes of [Ca^2+^]_i_ transients of female (magenta) and male (blue) myocytes with Mode 1 **(G)** and Mode 2 **(H)** ATP dynamics under 1-2 Hz stimulation. Data are presented as mean ± SD (female: N = 3 mice, *n* = 15 and 8 cells of Mode 1 and 2 sites, respectively; male: N = 3 mice, *n* = 16 and 15 cells of Mode 1 and 2 sites, respectively). All significant values are provided from a paired *t* test. Light symbols: individual cells; dark symbols: individual animals.

2 Hz pacing increased the amplitude of Mode 1 ATP gains in both sexes (**Figure 12C**) (females, 548 ± 31 µM → 722 ± 57 µM; males, 587 ± 58 µM → 811 ± 111 µM; each *P* < 0.0001), with males showing the larger absolute rise. Expressed as Δ[ATP]_i_, values in male myocytes again exceeded those in female myocytes at both 1 Hz (130 ± 58 µM vs. 91 ± 31 µM; P = 0.0140) and 2 Hz (173 ± 111 µM vs. 114 ± 57 µM; *P =* 0.0350) (**Figure 12E**).

Mode 2 regions, which exhibit ATP dips, behaved similarly at 1 Hz in both sexes. Raising the rate to 2 Hz shifted the dips to higher absolute concentrations (**Figure 12D**) (females: 380 ± 48 µM → 542 ± 67 µM, P = 0.0003; males: 383 ± 51 µM → 459 ± 73 µM; *P* = 0.0004), yet the depth of the dip (Δ[ATP]_i_) became markedly larger in males (−180 ± 73 µM) than in females (−66 ± 67 µM; *P* = 0.0009) (**Figure 12F**).

Notably, the accompanying [Ca^2+^]_i_ transients increased from 1 Hz to 2 Hz in both sexes in Mode 1 (**Figure 12G**) (female: 2.17 ± 0.43 → 2.74 ± 0.74 F/F_0_, male: 1.91 ± 0.32 → 2.34 ± 0.51 F/F_0_, each *P* < 0.0001) and Mode 2 regions (**Figure 12H**) (female: 2.29 ± 0.76 → 2.82 ± 1.04 F/F_0_, *P* = 0.0019; male: 2.06 ± 0.39 → 2.59 ± 0.60 F/F_0_, *P* < 0.0001), indicating that the frequency-dependent amplification of ATP gains and deepening of ATP dips occurs in the context of a larger Ca^2+^ signal. Taken together with our evidence of greater mitochondrial volume in males and tighter SR–mitochondrial coupling in females, these data support a model in which female myocytes meet heightened demand through architectural precision and efficient local production, whereas male myocytes rely on a mass-based scaling strategy that permits larger beat-coupled ATP swings but may approach capacity sooner. Thus, beat-synchronized ATP synthesis is modular and plastic, and its quantitative tuning differs between male and female cardiomyocytes.

## Discussion

In this study, we demonstrate that ATP is not continuously produced in adult ventricular myocytes but instead is generated in a beat-to-beat fashion through an on-demand process that is tightly coupled to EC coupling. We directly visualize rhythmic mitochondrial ATP transients with millisecond precision and show that they are phase-locked to each action potential and propagate to the cytosol. These transients appear as spatially confined microdomains—some showing ATP gains, others ATP dips—with similar amplitudes but different regional prevalence in male and female myocytes. Thus, ATP supply is organized into discrete, temporally synchronized bursts that match the periodicity of cellular demand, such that ventricular myocytes “live paycheck-to-paycheck,” producing just enough ATP during each beat to fuel contraction.

By resolving ATP within the mitochondrial matrix, we extend our prior findings of cytosolic ATP oscillations during EC coupling (Rhana *et al*., 2024) and identify their origin as upstream pulses of mitochondrial ATP production. The coexistence of gain and dip domains within the same cell, together with their regional segregation, underscores that mitochondrial output is both phasic and spatially modular. Pharmacological dissection reinforces a causal chain from Ca^2+^ release to matrix ATP pulses and cytosolic ATP changes: Ru360, thapsigargin, and FCCP abolish both ATP transients, establishing dependence on SR Ca^2+^ release and intact oxidative phosphorylation, whereas the MCU blocker Ru360 markedly attenuates beat-locked mito-iATP signals while preserving cytosolic Ca^2+^ transients, indicating that Ca^2+^ entry through the MCU is necessary to drive these ATP pulses. Blocking the adenine nucleotide translocator (ANT) with bongkrekic acid (BKA) reduced to near baseline mito-iATP transients while preserving Ca^2+^ cycling, demonstrating that beat-locked ATP synthesis and export through ANT are obligate steps in propagating mitochondrial ATP to the cytosol. Together with our earlier observation that Mode-1 cytosolic ATP gains are associated with increases in matrix Ca^2+^, whereas Mode-2 dips accompany Ca^2+^ loss (Rhana *et al*., 2024), these data support a unified mechanism in which SR-to-mitochondria Ca^2+^ transfer gates oxidative phosphorylation, ANT couples synthesis to export, and the resulting ATP pulses are read out as cytosolic ATP microdomains.

Mitochondrial bioenergetics and Ca^2+^-ATP coupling are highly conserved across mammalian species, and the fundamental processes we interrogate—mitochondrial Ca^2+^ uptake, tricarboxylic-acid (TCA) cycle activation, and ATP synthesis—operate via the same molecular machinery in mouse and human cardiomyocytes. Our use of 1 Hz stimulation allowed us to study these conserved mechanisms under conditions directly relevant to human resting heart rates, while taking advantage of the genetic tractability of the mouse for mechanistic perturbation. Thus, although the experiments were performed in mouse ventricular myocytes, the underlying principles of beat-resolved mitochondrial ATP supply are likely to generalize to human myocardium.

An interesting aspect of our data is that mito-iATP signals—presumably arising from intrafibrillar mitochondrial “strings”—extend over comparable distances in male and female myocytes (∼30 μm), implying that each string contains roughly the same number of individual mitochondria (∼15–20, assuming 1.5–2.0 μm length per mitochondrion in adult cells). Yet female myocytes have a smaller overall mitochondrial volume, and therefore fewer such strings per cell. On this reduced network, we find a higher per-mitochondrion density of the tethering protein mitofusin-2 (Mfn2), the catalytic α-subunit of the F_1_F_o_-ATP synthase (ATP5A), and, importantly, greater local overlap between VDAC and SERCA immunosignals, quantified by pixel-wise multiplication of binarized VDAC and SERCA masks and measurement of the area of overlap. These structural data indicate that, in females, each mitochondrion is more likely to be anchored close to SR Ca^2+^ release sites and endowed with higher ATP synthase content, providing two complementary advantages: tighter Ca^2+^ microdomains and higher catalytic capacity at each favored organelle.

This dual enrichment naturally explains the pronounced “hot spots” of ATP synthesis detected in female cells—zones where Ca^2+^ delivery (via increased VDAC–SERCA colocalization and higher Mfn2) and ATP synthase density coincide—as well as the relatively quiescent neighboring regions with fewer tethers and lower ATP5A. In contrast, male myocytes distribute a larger overall mitochondrial mass with lower per-mitochondrion ATP5A and less VDAC–SERCA overlap, achieving more spatially homogeneous rises in [ATP]_mito_ and relying on quantity rather than precision for energetic scaling. Ultrastructural evidence from other systems shows a denser cristae and heightened Ca^2+^ responsiveness in female cardiomyocyte mitochondria, together with systems-level data on female–male differences in mitochondrial function, support this view that architectural precision, coupled with enhanced catalytic machinery, allows female myocytes to meet localized energy demands efficiently. This sex-specific balance between mitochondrial abundance and per-organelle capacity may contribute to the greater resistance of female hearts to acute energetic stress and provides a structural framework for how modest changes in protein density and organelle positioning can reshape bioenergetic landscapes.

Consistent with this microdomain view, recent systems-genetics work by Cao *et al*. (2022) uncovered a sex-biased gradient in global mitochondrial supply and performance. Across >100 inbred mouse strains and two human heart cohorts, that study reported that female ventricles carry ∼10–20% fewer mitochondrial genomes and display coordinated down-regulation of nuclear- and mtDNA-encoded OXPHOS transcripts, and that mitochondria isolated from female hearts exhibit lower state-3 and maximal respiration and reduced complex II/IV flux. Moreover, gonadectomy plus hormone-replacement experiments showed that mitochondrial biogenesis is supported by androgens and restrained by estrogens, and overexpression of the male-biased fatty-acyl–CoA ligase ACSL6 improved both mitochondrial respiration and diastolic function, particularly in females. Viewed alongside our data, these findings suggest that female hearts rely on a lean but precisely deployed mitochondrial reserve that suffices at baseline yet may have limited headroom under sustained high workloads, whereas male hearts buffer demand with greater mitochondrial mass and more globally distributed OXPHOS capacity.

This architectural divergence has clear functional implications under stress. When pacing was raised from 1 to 2 Hz, mitochondrial ATP output reached the cytosol within one beat and preserved the microdomain pattern, but the way each sex accommodated the increased workload differed. Mode-1 (ATP-gain) amplitudes increased in both sexes, with larger absolute gains in males emerging only at higher frequency, pointing to recruitment of their larger mitochondrial pool. In Mode-2 (ATP-dip) regions, trough [ATP]_i_ rose in both sexes, but male myocytes exhibited deeper negative excursions, whereas female myocytes showed shallower dips despite preserved gains. These observations suggest that male myocytes meet demand primarily by scaling flux through their greater mitochondrial mass, tolerating larger beat-coupled swings in ATP at the cost of smaller net gains under high consumption, whereas female myocytes rely on tighter SR–mitochondrial coupling and high-capacity microdomains to limit the depth of local ATP deficits and match production to use within each hotspot.

Temporal analysis revealed additional sex-specific distinctions. Most female myocytes re-established diastolic ATP levels within ∼3.7 minutes after the onset of higher workload, consistent with rapid oxidative mobilization in pre-engaged, highly wired mitochondrial domains. Male myocytes exhibited a slower (∼4.5 minutes) but more gradual increase in ATP, aligning with recruitment of a larger, more heterogeneous mitochondrial pool that requires time to build up matrix NADH and activate latent oxidative phosphorylation capacity. We interpret the fast kinetic phase as activation of mitochondria already positioned near SR release sites and operating close to maximal dehydrogenase activity, and the slower phase as reflecting mobilization of more distant or less well-tethered domains that depend on cumulative Ca^2+^ and redox signals. This refinement of the classical concept of “mitochondrial reserve capacity” (Bertero & Maack, 2018) links reserve not only to enzyme excess but to spatial architecture and Ca^2+^ access, integrating structural sex differences (Cao *et al*., 2022) with functional disparities in ATP dynamics.

More broadly, our findings reframe cardiac energetics as modular and dynamic rather than homogeneous. Instead of a single, well-mixed ATP pool, mitochondrial output is organized into spatially restricted domains that deliver ATP in synchrony with each heartbeat, providing local control over supply and demand for ion transport, Ca^2+^ handling, and contraction. Biological sex remodels this topology through differences in mitochondrial abundance, tether density, and SR contact geometry, thereby shaping how the heart adapts to stress and matches energy supply with demand at the level of individual microdomains.

These principles resonate with observations in other excitable cells. In neurons, presynaptic mitochondria positioned near active zones supply ATP in proportion to firing demand, and disruption of this local coupling impairs synaptic transmission and plasticity (Rangaraju *et al*., 2014). In sinoatrial node pacemaker cells, earlier work showed that Ca^2+^-driven cAMP/PKA signaling scales mitochondrial ATP production to match changes in spontaneous firing on a seconds-to-minutes timescale (Yaniv *et al*., 2013), and our recent study in intact mouse sinoatrial node and isolated myocytes now demonstrates that ATP supply in this tissue is also organized into beat-locked cytosolic and mitochondrial ATP microdomains with high- and low-gain phenotypes, mapping a Ca^2+^-timed energetic hierarchy across the node (Munoz *et al*., 2025). Taken together with the present ventricular data, the convergence of spatially targeted mitochondria, activity-dependent Ca^2+^ entry, and localized, phasic ATP delivery suggests that modular, time-locked ATP supply is a conserved strategy for sustaining the function of excitable cells.

Although we did not perform a full calibration of absolute matrix [ATP], the robust mito-iATP oscillations we observe have important implications for mitochondrial ATP levels. Based on the apparent K_d_ of ∼0.5 mM for iATPSnFR2 (Marvin *et al*., 2024), fluorescent changes of the magnitude we detect are most consistent with free mitochondrial [ATP] in the low-millimolar range (∼1–5 mM), rather than the ∼7.5–8 mM values inferred from biochemical or NMR measurements of total ATP. Taken together with our prior finding that cytosolic [ATP]i is <1 mM (Rhana *et al*., 2024), this yields an apparent paradox: if neither compartment harbors very high free ATP, how do we reconcile these values with the ∼8 mM total cellular ATP reported by classic biochemical and NMR studies (e.g., Schwenke *et al*. (1981))? A plausible resolution, aligned with recent conceptual work by Eisner and Murphy (2024), is that most cellular ATP is bound to proteins or sequestered in compartments that are largely invisible to genetically encoded sensors, which report free nucleotide. In this revised view, both matrix and cytosol operate with relatively low free ATP, explaining why inhibition of oxidative phosphorylation does not lead to the large cytosolic ATP increases predicted by earlier models and why modest fractional changes in ATP can exert substantial control over ATP-dependent enzymes and channels. Motion artifacts are unlikely to account for our signals, as a JFX554 HaloTag reference channel showed no beat-locked changes while the mito-iATP ratio tracked linearly with non-ratiometric mito-iATP fluorescence, indicating that the ratio readout faithfully reports changes in matrix ATP rather than movement of mitochondria within the confocal volume.

Our findings also sharpen the link between cardiac energetics and coronary blood flow, placing functional hyperemia at the center of beat-resolved ATP homeostasis. Classic work reviewed by Feigl (1983) established that coronary flow is dynamically regulated to match myocardial oxygen demand, with local metabolic signals acting in tandem with myogenic and neural inputs to tune vascular resistance. Recent studies extend this framework by highlighting microvascular rarefaction and altered capillary–myocyte coupling as key determinants of contractile reserve and arrhythmia susceptibility (Manning *et al*., 2024; Manning *et al*., 2025). Within this context, the rapid, beat-synchronized nature of mitochondrial ATP synthesis that we describe implies that continuous delivery of oxygen and substrates through the coronary microcirculation is essential not just for long-term metabolic integrity but for preserving the energetic fidelity of each heartbeat. Even brief disruptions of blood flow or impaired functional hyperemia could compromise the mitochondrial response to [Ca^2+^]_i_ transients, degrading contractile performance and electrical stability within a few cycles. Thus, perfusion and metabolism emerge as moment-to-moment co-regulators of cardiac function, tightly integrated through beat-locked ATP microdomains.

The CMIC axis proposed by Santana and Earley (In press) provides a useful lens for interpreting these findings across scales: microvascular delivery sets boundary conditions for oxidative metabolism, mitochondria generate ATP in a structured network, and ATP-dependent membrane/SR effectors convert local ATP availability into excitability and Ca²⁺ handling. While we did not directly manipulate the capillary network here, by resolving beat-locked matrix ATP microdomains and their propagation to cytosol—together with MCU and ANT dependence—we establish the mitochondrial “middle link” of CMIC on the timescale of individual heartbeats. This framing predicts that limitations in oxygen/substrate delivery or capillary–myocyte coupling will be expressed as reduced fidelity of beat-synchronized ATP supply, with rapid consequences for electrical and mechanical stability. In addition, our male–female comparisons suggest that CMIC tuning can be achieved by distinct design strategies: greater mitochondrial mass in males versus tighter SR–mitochondrial wiring and higher per-organelle catalytic capacity in females, producing different frequency-dependent scaling of diastolic versus beat-locked ATP components under workload stress.

In summary, ATP supply in adult ventricular myocytes is shaped by a network of beat-coupled, spatially organized production units. Mitochondria discharge ATP in phasic bursts that are aligned with each action potential, exported via ANT, and propagated to the cytosol to support rhythmic contractile demands. Sex-specific differences in mitochondrial mass, tethering, and SR contact geometry tune this architecture, with male myocytes leveraging mitochondrial abundance and female myocytes achieving stability through precision wiring and higher per-organelle capacity. By providing the first direct visualization of beat-to-beat mitochondrial ATP transients in adult heart cells, our work compels a revision of traditional bioenergetic models—from static, well-mixed ATP pools to dynamic, microdomain-based supply—and suggests that cardiac metabolic resilience is governed as much by spatial design and Ca^2+^ access as by absolute mitochondrial capacity.

## First author profile

Paula Rhana, PhD, is an AHA Postdoctoral Fellow in Physiology & Membrane Biology, UC Davis. Trained in Biomedicine (BSc, FUMEC, Brazil) and Biochemistry & Immunology (MSc, PhD, UFMG, Brazil), Dr. Rhana discovered how voltage-gated Na^+^ and Ca^2+^ channels drive breast tumor progression. Upon joining the Santana lab in 2022, she led the first effort to identify and characterize beat-to-beat mitochondrial ATP transients in adult cardiomyocytes using combined imaging and electrophysiological approaches. Her current research centers on mapping sex-specific metabolic microdomains that tune cardiac excitability, with the ultimate goal of translating these energetic insights and fluorescent-sensor innovations into new strategies for preventing and treating heart failure.

## Abbreviations

SR: sarcoplasmic reticulum
VDAC: voltage-dependent anion channel
MCU: mitochondrial Ca^2+^ uniporter
ANT: adenine nucleotide translocase.

## Data availability statement

The raw data files supporting all findings presented in this paper are available from the corresponding author.

## Author contributions

PR and LFS conceived and designed the work. PR and CM acquired and analyzed data for the paper. PR, CM, and LFS wrote the paper. All authors have approved the final version of the manuscript and agreed to be accountable for all aspects of the work. All persons designated as authors qualify for authorship, and all those who qualify for authorship are listed.

## Acknowledgements

We thank Mr. Josh Tulman for original illustrations. We also thank Dr. Declan Manning and Dr. Manuel F. Muñoz for reading the manuscript.

## Figures

**SI Figure 1.**
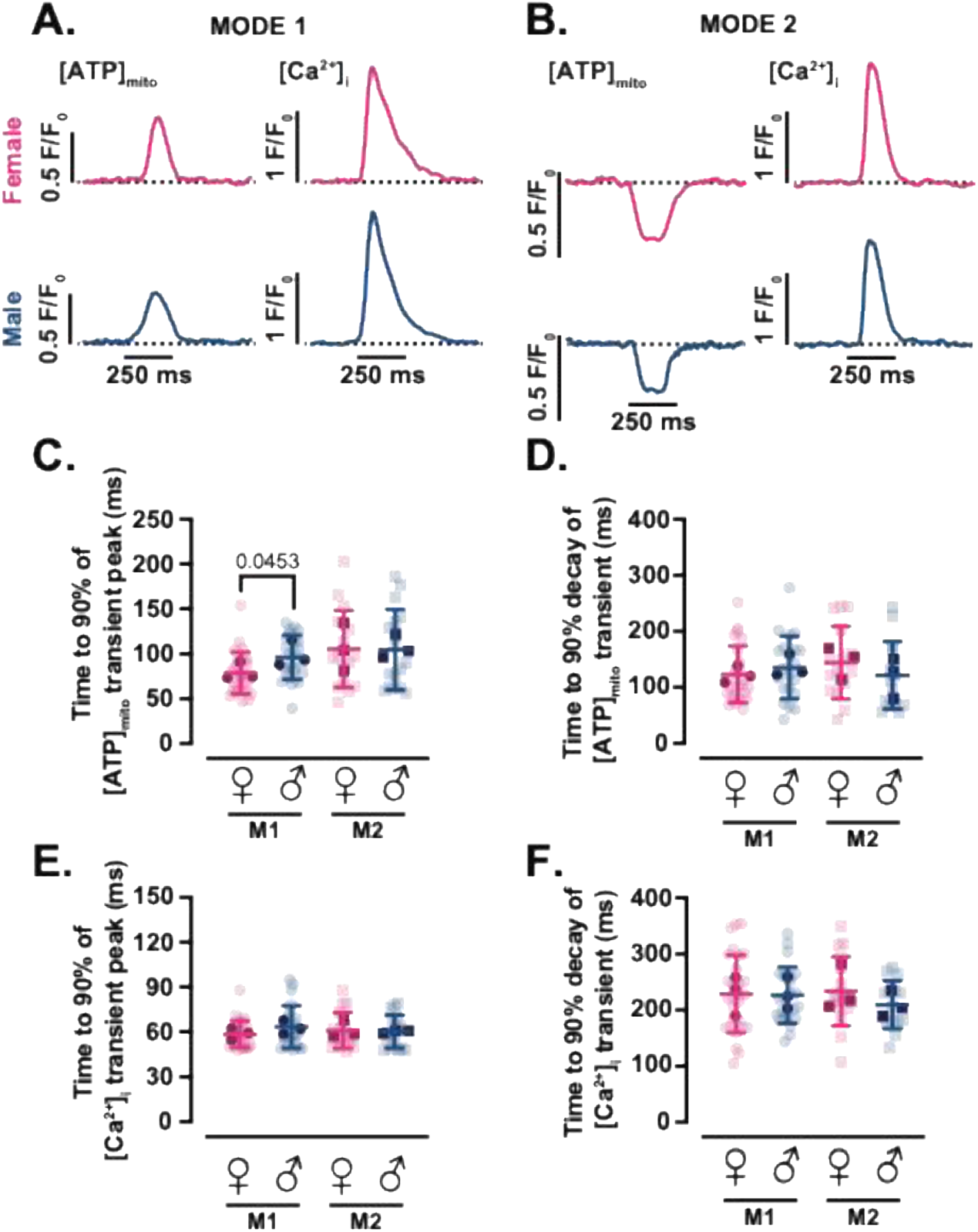
Kinetics of mitochondrial ATP and [Ca²⁺]_i_ transients. Representative time course from line scan images of [ATP]_mito_ (left) and [Ca^2+^]_i_ (right) in female (magenta) and male (blue) myocytes with Mode 1 **(A)** or Mode 2 **(B)** ATP dynamics. Scatter plots for time to 90% of the [ATP]_mito_ transient peak **(C)** and decay **(D)** in Mode 1 and Mode 2 regions of female (magenta) and male (blue) myocytes. Scatter plots for time to 90% of the [Ca^2+^]_i_ transient peak **(E)** and decay **(F)** in Mode 1 and Mode 2 regions of female (magenta) and male (blue) myocytes. Data are presented as mean ± SD (female: N = 3 mice, *n* = 28 and 17 cells of Mode 1 and 2 sites, respectively; male: N = 3 mice, *n* = 24 and 16 cells of Mode 1 and 2 sites, respectively). All significant values are provided from a nested *t* test. Light symbols: individual cells; dark symbols: individual animals.

**SI Figure 2.**
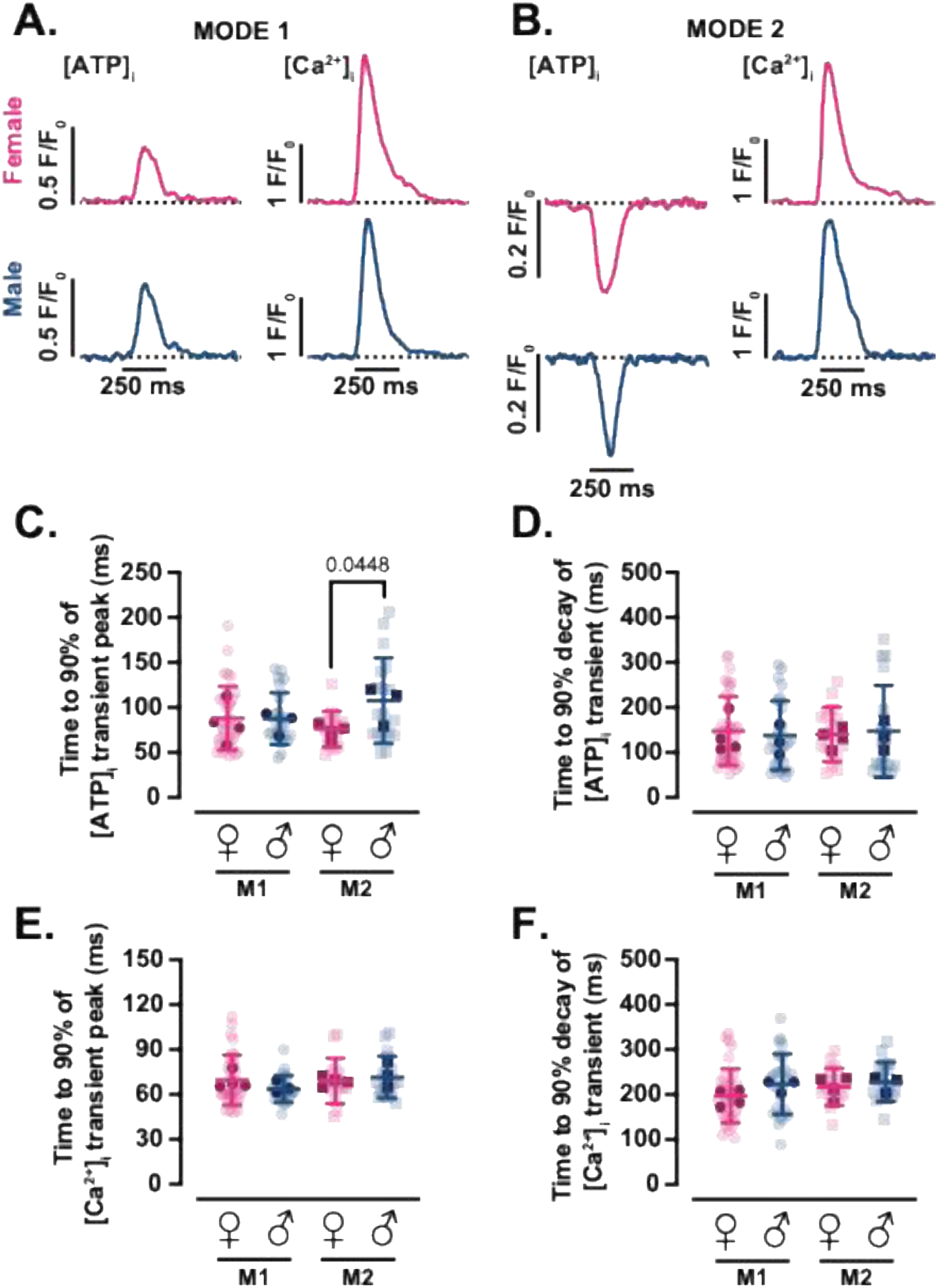
Kinetics of cytosolic ATP transients and [Ca²⁺]_i_ transients. Representative time course from line scan images of [ATP]_i_ (left) and [Ca^2+^]_i_ (right) in female (magenta) and male (blue) myocytes with Mode 1 **(A)** or Mode 2 **(B)** ATP dynamics. Scatter plots for time to 90% of the [ATP]_i_ transient peak **(C)** and decay **(D)** in Mode 1 and Mode 2 regions of female (magenta) and male (blue) myocytes. Scatter plots for time to 90% of the [Ca^2+^]_i_ transient peak **(E)** and decay **(F)** in Mode 1 and Mode 2 regions of female (magenta) and male (blue) myocytes. Data are presented as mean ± SD (female: N = 4 mice, *n* = 34 and 15 cells of Mode 1 and 2 sites, respectively; male: N = 3 mice, *n* = 24 and 18 cells of Mode 1 and 2 sites, respectively). All significant values are provided from a nested *t* test. Light symbols: individual cells; dark symbols: individual animals.

